# All-optical closed-loop voltage clamp for precise control of muscles and neurons in live animals

**DOI:** 10.1101/2022.06.03.494532

**Authors:** Amelie C.F. Bergs, Jana F. Liewald, Silvia Rodriguez-Rozada, Qiang Liu, Christin Wirt, Artur Bessel, Nadja Zeitzschel, Hilal Durmaz, Adrianna Nozownik, Maëlle Jospin, Johannes Vierock, Cornelia I. Bargmann, Peter Hegemann, J. Simon Wiegert, Alexander Gottschalk

## Abstract

Excitable cells can be stimulated or inhibited by optogenetics. Since optogenetic actuation regimes are often static, neurons and circuits can quickly adapt, allowing perturbation, but not true control. Hence, we established an optogenetic voltage-clamp (OVC). The voltage-indicator QuasAr2 provides information for fast, closed-loop optical feedback to the bidirectional optogenetic actuator BiPOLES. Voltage-dependent fluorescence is held within tight margins, thus clamping the cell to distinct potentials. We established the OVC in muscles and neurons of *Caenorhabditis elegans*, and transferred it to rat hippocampal neurons in slice culture. Fluorescence signals were calibrated to electrically measured potentials, and wavelengths to currents, enabling to determine optical I/V-relationships. The OVC reports on homeostatically altered cellular physiology in mutants and on Ca^2+^-channel properties, and can dynamically clamp spiking. Combining non-invasive imaging with control capabilities of electrophysiology, the OVC facilitates high-throughput, contact-less electrophysiology in individual cells and paves the way for true optogenetic control in behaving animals.

## Introduction

Identifying connections between distinct neurons and their contribution to driving behavior is a central issue in neuroscience^1,2^. To explore such relations, methods to record and concurrently regulate neural activity are needed^3^. Also, high-throughput screening approaches in excitable cell physiology require application of such methods. Diverse approaches to control or observe excitable cell function are in use. Patch-clamp electrophysiology provides superior temporal accuracy and sensitivity, but is limited by its invasiveness^4-6^. Ca^2+^-imaging, applicable in intact living organisms, enables integrating cell physiology and behavioral output^7-9^. However, due to sensor Ca^2+^-buffering and the non-linear correlation of cytosolic Ca^2+^-concentration and membrane voltage, this technique suffers from comparably low temporal resolution, and fails to resolve subthreshold voltage transients or high-frequency action potentials (APs). Further, since Ca^2+^-concentration does not fall below basal cytosolic levels in most neurons, Ca^2+^-imaging is unsuited to reveal synaptic inhibition. Instead, membrane potential can be imaged *via* <u>g</u>enetically <u>e</u>ncoded <u>v</u>oltage indicators (GEVIs), e.g. rhodopsin-based GEVIs, among others^10-17^. The fluorescence of retinal, embedded in rhodopsins, reliably monitors voltage dynamics at millisecond timescales. Since rhodopsin-based GEVIs emit near-infrared light, they can be multiplexed with optogenetic actuators of membrane currents, to selectively photostimulate or inhibit activity of excitable cells with high spatiotemporal precision^18-20^. In the “(i)-Optopatch” approach, the blue-light activated channelrhodopsin (ChR) CheRiff was used together with the voltage indicator QuasAr2, allowing to unidirectionally steer and observe, but not to fully control neuronal activity^11,12,21^. An optical dynamic clamp (ODC) used archaerhodopsin to mimic K^+^-currents, absent in immature cardiomyocytes, in closed loop with electrophysiological feedback^22^. Bidirectional optical modulation and readout of voltage was implemented as light-induced electrophysiology (LiEp) for drug screening, however, without a feedback loop^23^. A bidirectional approach with feedback – “optoclamp” – used ChR2 and halorhodopsin (NpHR) as actuators and extracellular microelectrode arrays instead of GEVIs^24^. This approach clamped average firing rates in neuronal ensembles, following indirect voltage readout.

True all-optical control over excitable cell activity with closed-loop feedback, as in voltage-clamp electrophysiology, should combine two opposing optogenetic actuators for de- and hyperpolarization, as well as a GEVI. Such a system could respond to intrinsic changes in membrane potential. It would also prevent inconsistent activity levels arising from cell-to-cell variation in optogenetic tool expression levels. These are usually not taken into account, particularly when static stimulation patterns are used, while a feedback system could increase stimulation until the desired activity is reached. The system should synergize the non-invasive character of imaging methods with the control capabilities of electrophysiology.

Here, we established an <u>o</u>ptogenetic <u>v</u>oltage-<u>c</u>lamp (OVC) in *Caenorhabditis elegans* and explored its use in rat hippocampal organotypic slices. The OVC uses QuasAr2 for voltage read-out^11,20^, and BiPOLES, a tandem protein comprising the depolarizer Chrimson and the hyperpolarizer *Gt*ACR2, stimulated by orange and blue light, respectively, for actuation^19,25-28^. Spectral separation and their balanced 1:1 expression in BiPOLES enabled gradual transitions from depolarized to hyperpolarized states, and *vice versa*. QuasAr2 fluorescence was used to compute a feedback of wavlength-adapted light signals transmitted to BiPOLES, in closed-loop at up to 100 Hz, thus keeping the voltage-dependent fluorescence at a desired level. We characterized the system in body-wall muscle cells (BWMs), as well as in cholinergic and GABAergic motor neurons. Simultaneous measurements allowed calibrating fluorescent signals to actual membrane voltages, and passively presented wavelength pulses to currents. In *unc-13* mutants, the OVC readily detected altered excitability of muscle, as a response to the reduced presynaptic input. In *egl-19* VGCC gain-of-function (g.o.f.) mutants, an optical I/V-relationship of the mutated channel could be obtained, comparing well to electrophysiological measurements. In rodent neurons, the OVC also modulated fluorescence and voltage, yet with a smaller range, due to the resting potential being close to Cl^-^-reversal potential. Last, in spontaneously active tissues, i.e., pharyngeal muscle and the motor neuron DVB^29-32^, the OVC could dynamically follow and counteract native APs, and suppress associated behaviors.

## Results

### Reading GEVI fluorescence to steer optogenetic actuators with light feedback in closed-loop

An OVC should measure voltage-dependent fluorescence of an excitable cell, e.g. *via* a rhodopsin-GEVI^10,11^, and provide adjusted light-feedback to optogenetic actuators of membrane voltage, e.g. cation- and anion-selective rhodopsin channels^19,28,33,34^ (**Fig. 1A**). The OVC needs to work in closed-loop, to quickly counteract intrinsic activity. We first implemented suitable hard- and software (**Fig. 1B**): To excite fluorescence of rhodopsin GEVIs (e.g. QuasAr2, emitting in the far-red)^10,11^, we expand a 637 nm laser to cover 0.025 mm². GEVI fluorescence of a region of interest (ROI) is monitored by a camera, and compared to a target value (**Fig. 1C, Extended data Fig. 1A-C**). Light feedback is sent to the sample from a monochromator, whose wavelength limits can be pre-selected to match the chosen hyperpolarizing and depolarizing optogenetic tools’ maximal activation, and that can adjust wavelength at 100 μs and 0.1 nm temporal and spectral resolution, respectively.

**Fig. 1.**
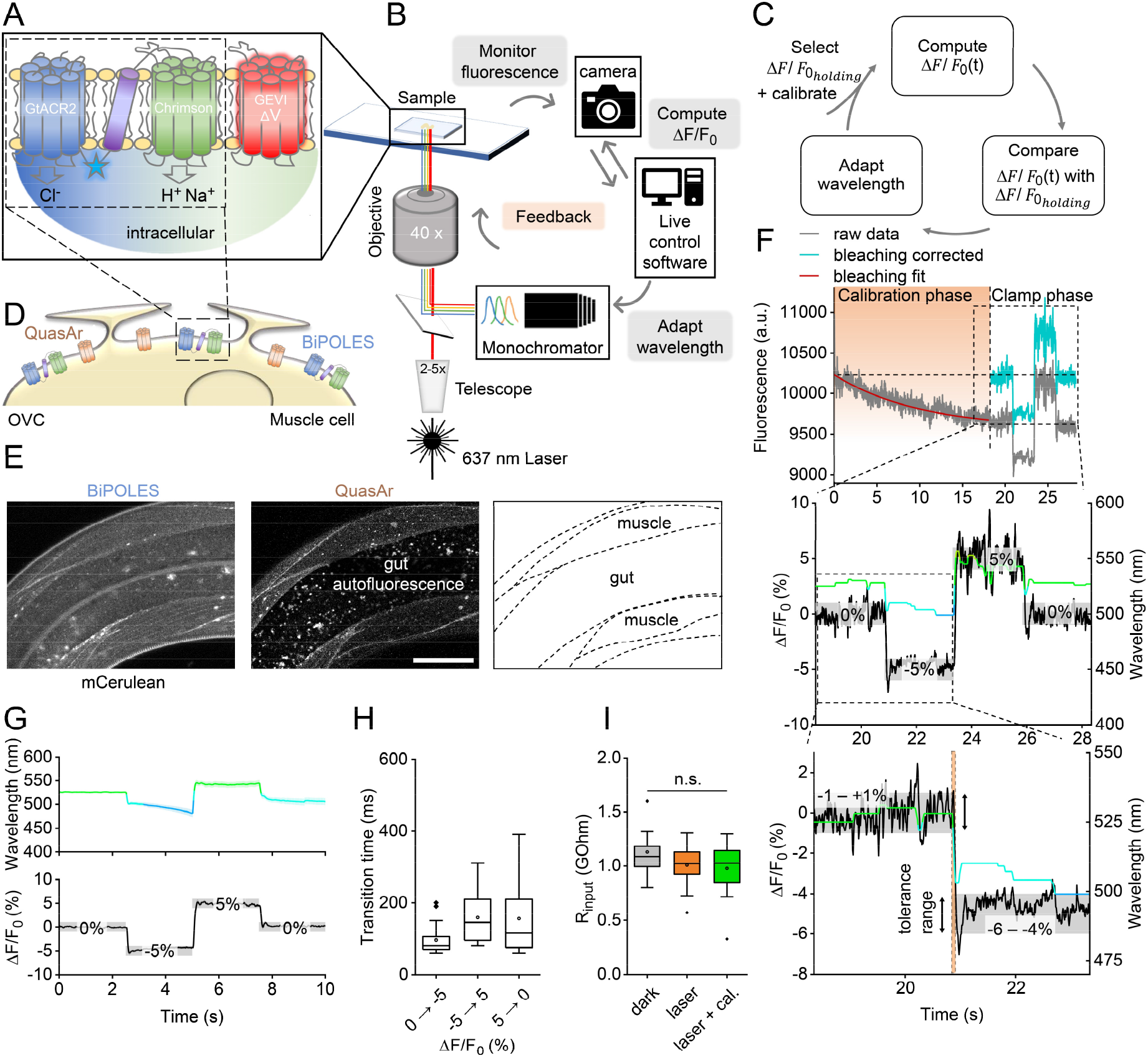
Components, setup and functionality of the OVC. **(A)** Molecular OVC components: Two counteracting optogenetic actuators for de- and hyperpolarization (GtACR2, Chrimson; ★: mCerulean), and a voltage indicator. **(B)** Hardware setup and communication: Membrane voltage is monitored *via* a fast and sensitive (sCMOS or EMCCD) camera; gray values are processed by live control software. Based on the difference to the set holding value, a feedback of light from a monochromator is sent to the optogenetic actuators. **(C)** OVC feedback mechanism: After selection of clamping parameters and calibration for bleaching correction, ΔF/F_0_ is calculated and compared with the set value, to adapt the wavelength accordingly, in closed-loop. **(D)** OVC (QuasAr2 and BiPOLES) in BWMs. **(E)** Confocal fluorescence z-projection. BiPOLES (mCerulean) and QuasAr2 in BWM membranes, scheme (dashed lines represent muscle plasma membranes). Scale bar, 50 μm. **(F)** OVC four-step protocol (0, -5, 5, and again 0 % ΔF/F_0_) in BWMs; insets and below: close-up. Wavelength shown in the respective color, holding values and tolerance range (grey boxes) are indicated for each step. Orange shade in lower panel: transition period to reach tolerance range. **(G)** Upper panel: Overlay of mean (± S.E.M.) wavelength and (lower panel) fluorescence traces (n=24; holding values: 0, -5, 5, 0 % ΔF/F_0_). **(H)** Times required for the indicated 5 and 10 % ΔF/F_0_ transitions. **(I)** Membrane resistance of BWM cells, before, and during 637 nm laser illumination (orange), and during laser + calibration light (green); n=12-13. One-way ANOVA with Bonferroni correction (laser / dark: p = 0.65; laser+cal. / dark: p = 0.39; laser+cal. / laser: p = 1). In (H and I), box plots (median, 25-75^th^ quartiles); open dot: mean; whiskers: 1.5x IQR.

Communication between camera and monochromator is provided by a custom-written script in Beanshell (part of μManager interface^35^), processing incoming gray values into relative changes of fluorescence (ΔF/F_0_) (**Fig. 1C, Extended data Fig. 1A-C**), and operating at ca. 100 fps. Since GEVIs exhibit photobleaching, ΔF/F_0_ values would gradually deviate from actual voltage levels. Thus, for each recording, an initial calibration phase is used to calculate correction parameters (**Extended data Fig. 1D, E**). Once the system can access bleach-corrected ΔF/F_0_ values, it feeds them into a decision tree algorithm (**Extended data Fig. 1A**), where they are compared to a desired holding ΔF/F_0_ value. Deviation between target and actual ΔF/F_0_ determines the wavelength change of the monochromator (I(ntegral)-controller). An alternative algorithm, using a PID controller^36^ and Kalman filter^37^ for sensor smoothing, did not increase overall system performance (see methods; **Extended data Fig. 2**).

### Combining single actuators with QuasAr2 for unidirectional steering of membrane voltage

First, we expressed QuasAr2 in *C. elegans* body wall muscles (BWMs). Expression levels were very uniform across and within strains, ensuring equal OVC activity in different genetic backgrounds (**Extended data Fig. 1F**). 20 s calibration under 637 nm laser light sufficed to estimate photobleaching parameters. 300 μW/mm^2^ blue light or presenting the full spectrum (400 to 600 nm) caused no additional bleaching and did not affect QuasAr2 fluorescence (**Extended data Fig. 1D, E, G**). Thus, monochromator light did not influence bleaching-corrected fluorescence and estimated membrane voltages. Initially, to test the feedback loop, we assessed different actuators and configurations. First, *Chlamydomonas reinhardtii* ChR2(H134R) or *Guillardia theta* anion channelrhodopsin *Gt*ACR2 were expressed in cholinergic motor neurons (**Extended data Fig. 3A**). This allowed manipulating muscle voltage indirectly via light-induced (de-)activation of motor neurons, and to adjust QuasAr2 fluorescence to values between +20 and -15 % ΔF/F_0_, respectively (**Extended data Fig. 3B-D**). However, fluorescence (i.e., voltage) returned to baseline only by intrinsic membrane potential relaxation (**Extended data Fig. 3D-G**), thus limiting temporal resolution: For ChR2, fluorescence reached the target range after 150 ± 12.7 ms and relaxed within 678 ± 148.9 ms (254 ± 20.4 ms and 239.5 ± 29.9 ms, respectively, for *Gt*ACR2). Next, we assessed combinations of spectrally distinct, opposing actuator pairs, like ChR2 (470 nm) and *Gt*ACR1 (515 nm), or *Natronomonas pharaonis* halorhodopsin NpHR (590 nm). Optogenetic de- or hyperpolarization of cholinergic motor neurons affects muscle activation and evokes body contraction or elongation^33,38^ (**Extended data Fig. 4A-D**). Yet, one actuator was typically outperformed by the other, impeding precise control of membrane potential. Likely, separate expression led to variable relative amounts of the tools. We thus resorted to 1:1 expression via BiPOLES.

### BiPOLES enables bidirectional voltage-clamping in *C. elegans* muscle

The tandem protein BiPOLES combines de- and hyperpolarizers Chrimson and GtACR2 (590 nm and 470 nm, respectively), linked as one sequence^27^ (**Fig. 1A**). BiPOLES activation with a 400-600 nm ramp evoked robust bidirectional effects on body length (**Extended data Fig. 4E**). We co-expressed BiPOLES and QuasAr2 in BWMs (**Fig. 1D, E**). Low levels of BiPOLES were found at the plasma membrane (PM) and in few intracellular aggregates, while QuasAr2 localized mainly to the PM. As QuasAr2 excitation (637 nm) causes some activation of Chrimson (their spectra overlap^26,39^; **Extended data Fig. 5A**), we needed to counteract its effects *via* GtACR2, by compensatory light from the monochromator. The wavelength was adjusted until -5 and 5 % ΔF/F_0_ could be maintained for ca. 5 s. Additional assays ensured that compensatory light reinstates normal function: In animals expressing the OVC in BWMs or cholinergic neurons, 637 nm laser light diminished motor behavior. However, compensatory light restored body length and swimming behavior (**Extended data Fig. 5B-E, Extended data Video 1**), and BWMs expressing the OVC showed voltage fluctuations comparable to animals expressing QuasAr2 only (**Extended data Fig. 5F-H**). 637 nm laser light did not fully activate Chrimson, as it could be further excited by 590 nm (300 μW/mm²) light, increasing QuasAr2 fluorescence by 2.3 % (**Extended data Fig. 5I**). After bleaching correction, prior to each individual experiment (R^2^ of the exponential fit was always >0.8, in most cases even >0.95; **Extended data Fig. 6A**), the OVC generated incremental changes in wavelength that closely followed the fluctuating fluorescence signals (i.e., membrane voltage; **Fig. 1F**), as soon as the tolerance range for holding ΔF/F_0_ was exceeded. Cells were reliably and quickly (**Extended data Fig. 6B, C**), clamped to holding values between -5 and 5 % ΔF/F_0_ (**Fig. 1F, G**). Due to the bidirectionality and live feedback, the BiPOLES-OVC acted significantly faster than when using single actuators, particularly for the return towards resting potential: Transition times were only 147.7 ± 25.5 ms (5→0 %) and 88.0 ± 5.7 ms (−5→0 %) for BiPOLES, compared to 678.6 ± 148.9 ms for ChR2 and 239.5 ± 29.9 ms for *Gt*ACR2 (Fig. 1H, Extended data Figs. 3G, 6C). The OVC allowed continuous bidirectional clamping for extended periods (**Extended data Fig. 6D-G**). Once the closed-loop control was interrupted, membrane voltage approached baseline and higher fluorescence fluctuations were observed (1.87 ± 0.1 vs. 0.78 ± 0.05 % ΔF/F_0_ during -5 % clamping; 1.89 ± 0.14 % ΔF/F_0_ for animals expressing only QuasAr; **Extended data Fig. 5F-H**). Importantly, the cell could also be actively steered back to the initial fluorescence level by the OVC, indicating that there is no hysteresis associated with the closed-loop approach (**Fig. 1F, G**).

### Calibrating QuasAr2 fluorescence and membrane potential in BWMs

To calibrate the OVC system and determine the actually accessible voltage range, we measured voltage and fluorescence simultaneously (**Extended data Fig. 7A**). Concurrent laser (637 nm) and compensation illumination (520 nm) did not significantly alter membrane resistance (**Fig. 1I**), or APs, that could be observed by fluorescence and patch-clamp simultaneously (**Fig. 2A; Extended data Fig. 7B, C**). The dual illumination also did not alter membrane potential; however, adding 470 nm light could hyperpolarize the cell by about 16 mV (**Extended data Fig. 7D**). Small voltage fluctuations, likely representing EPSPs, were similar in dark and light conditions by amplitude and frequency (**Extended data Fig. 7E-G**). Thus, BiPOLES activation, despite the open channels, did not lead to general shunting of membrane potential in *C. elegans* muscle.

**Fig. 2.**
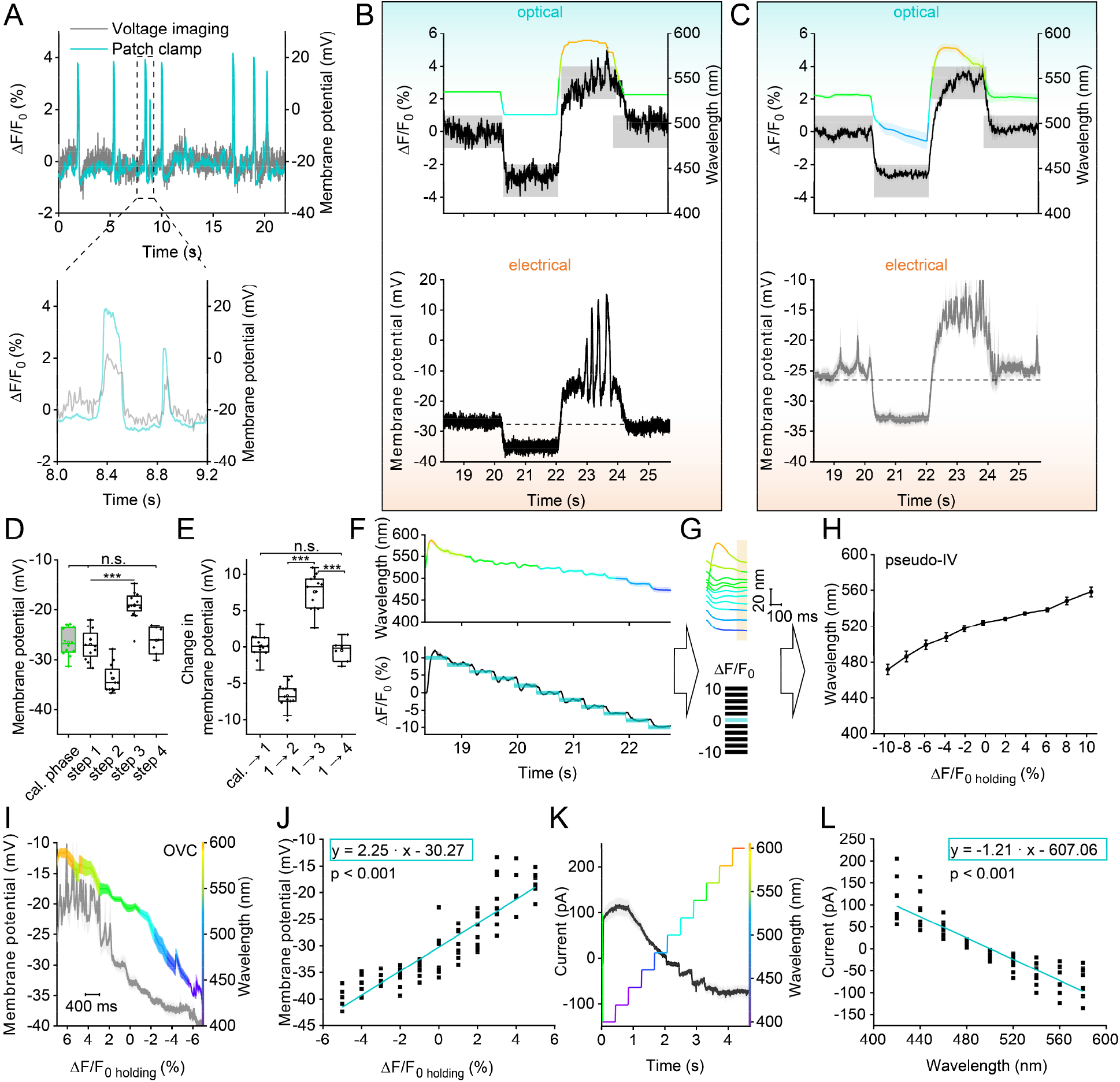
Bi-directional optical clamping and calibration of membrane voltage and currents in BWMs. **(A)** APs in simultaneous patch-clamp and fluorescence recordings during OVC calibration phase (637 nm laser + calibration wavelength). Inset for close-up. **(B)** Simultaneous patch-clamp and fluorescence recording of an OVC four-step protocol (0, -3, 3, 0 % ΔF/F_0_). Upper panel: Fluorescence recording with indicated wavelength adaptation and tolerance ranges. Lower panel: Corresponding simultaneous whole-cell patch-clamp voltage recording. **(C)** Overlay of mean (± S.E.M.) wavelength, fluorescence (upper panel) and voltage traces (lower panel; n=12) as in (B). **(D)** Statistical analysis of data in (C); absolute voltages of transition events during simultaneous OVC measurements (cal / step 1: p = 1, cal / step 4: p = 1, step 1 / step 4: p = 1, step 1 / step 2: p = 4.1E-6, step 1 / step 3: p = 1.6E-7, step 2 / step 3: p = 4.5E-16). **(E)** Voltage modulation for the respective steps of the OVC protocol in (D) (cal → step1 / step 1 → step 2: p = 7.51E-11, step 1 → step2 / step1 → step3: p = 1.04E-21, step1 → step3 / step1 → step4: p = 1.13E-10, cal → step1 / step1 → step4: p = 1). **(F)** Distinct ΔF/F_0_ steps were presented by the OVC (lower panel: blue shades, tolerance ranges; black trace, actual ΔF/F_0_ mean ± S.E.M.), and the required wavelengths recorded (mean ± S.E.M. nm; upper panel). **(G, H)** Determining the wavelengths (≙ currents) at the end (highlighted in orange) of each ΔF/F_0_ clamp step (≙ voltage) to deduce a pseudo-I/V relation (n=19). **(I, J)** Simultaneous OVC / electrophysiology experiment, setting ΔF/F_0_ clamp steps, measuring resulting membrane potentials, to determine a calibration regression (membrane potential over ΔF/F_0_, n=5-9). **(K, L)** Voltage clamp (at -24 mV) measurement of currents resulting from wavelength steps, to determine a calibration regression (current over wavelength, n=8). One-way ANOVA with Bonferroni correction. ***P ≤ 0.001 in (D, E), box plots (median, 25-75^th^ quartiles); open dot: mean; whiskers: 1.5x IQR.

In simultaneous electrophysiology and optical OVC experiments, induced fluorescence changes (−3 to +3 % ΔF/F_0_, and returning back to 0% ΔF/F_0_) modulated voltage by ca. -7 to +8 mV (**Fig. 2B-E**), i.e. ca. 15 mV range per 6 % ΔF/F_0_ (24 mV per 10 % ΔF/F_0_; given the linear fluorescence-voltage relation of QuasAr2; Ref.11). Note, a smaller ΔF/F_0_ range was chosen to facilitate patch-clamp measurements; briefer optical clamp periods (400 ms instead of 2 s) allowed accessing broader voltage ranges (see below). We verified the fidelity of the OVC by assessing whether the initially deduced calibration function may cause progressive errors during the voltage clamp phase. Correlating induced ΔF/F_0_ traces to evoked membrane potentials revealed no (increasing) deviation of actual voltages from OVC-imposed trajectories (**Extended data Fig. 7H-K**).

### Measuring all-optical I/V relationships

The OVC allowed reliable optical voltage clamping. We wondered if we could also use it to acquire all-optical I/V-relationships. To this end, we devised a different software “pseudo I/V-curve” (**Extended data Fig. 8A**), which can consecutively present different ΔF/F_0_ clamp steps. First, we performed optical experiments, relating different ΔF/F_0_ clamp values (= voltage equivalent) to observed wavelengths (= current equivalent), required to achieve the respective fluorescence steps (**Fig. 2F, G**). These purely optical experiments (in intact animals) showed that a range of at least ±10 % ΔF/F_0_ can be achieved with wavelengths of ca. 470-585 nm, different to the simultaneous OVC/patch-clamp experiments (**Fig. 2H; Extended data Fig. 8B**). As an inverse control, we presented wavelengths and achieved a congruent output ΔF/F_0_ level (**Extended data Fig. 8C, D**). Next, we calibrated the ΔF/F_0_ steps to actual voltages, and the wavelengths to actual currents. We again carried out simultaneous OVC/patch-clamp experiments and examined membrane voltage as a function of the pre-set clamp fluorescence (**Fig. 2I**). Individual steps were shortened to 400 ms, which allowed covering a range of ±5 % ΔF/F_0_. The voltages determined and the range covered corresponded to those of the ±3 % ΔF/F_0_ measurements using the ‘standard’ OVC protocol, i.e. ca. 22 mV (−40 to -18 mV) for ±5 % ΔF/F_0_ (**Fig. 2J**). Likewise, we determined membrane currents (at -24 mV, i.e. BWM resting potential), induced by step changes in applied wavelengths (**Fig. 2K**). BiPOLES mediated currents in a total range of ∼190 pA, running almost linearly between 420 and 580 nm (**Fig. 2L**). Linear regression estimations allowed relating membrane voltages to ΔF/F_0_ and currents to wavelengths, and to identify calibration parameters (**Fig. 2J, L**; methods). Significance levels for the coefficients were <0.001, indicating high precision. Thus, averaged optical data can be converted to voltage and currents.

### Demonstrating homeostatic changes in muscle excitability using the OVC

We explored the utility of the OVC to assess divergent cell physiology. *unc-13(n2813)* mutants lack an essential synaptic vesicle priming factor^40^ and thus exhibit largely reduced postsynaptic currents upon ChR2-stimulation of motor neurons^41^. Yet, *unc-13* and other neurotransmission mutants showed enhanced muscle contraction when BWMs were directly photostimulated. We hypothesized that muscles homeostatically change their excitability. To explore if the OVC can detect this, we induced a +5 % ΔF/F_0_ depolarization step: In *unc-13(n2813)* mutants, the OVC required significantly blue-shifted light (i.e., less Chrimson activation) to induce the same level of depolarization as in wild type (534.2 ± 2.8 vs. 550.4 ± 3.1 nm; **Fig. 3A, B**). Muscles in *unc-13* animals may exhibit higher excitability to balance the lower excitatory input they receive. Though no significant change in membrane resistance was observed (**Fig. 3C**), the amplitude of induced voltage increases was higher in *unc-13* mutants when we injected current ramps (**Fig. 3D-F**). Similarly, APs measured by voltage imaging were significantly increased by amplitude and duration (**Fig. 3G, H, J**). Thus, in response to a lack of presynaptic input, ion channels shaping muscle APs, i.e. voltage-gated Ca^2+^ (EGL-19) and K^+^-channels (SLO-2, SHK-1)^42,43^, may undergo homeostatic changes to enable the observed higher muscle excitability.

**Fig. 3.**
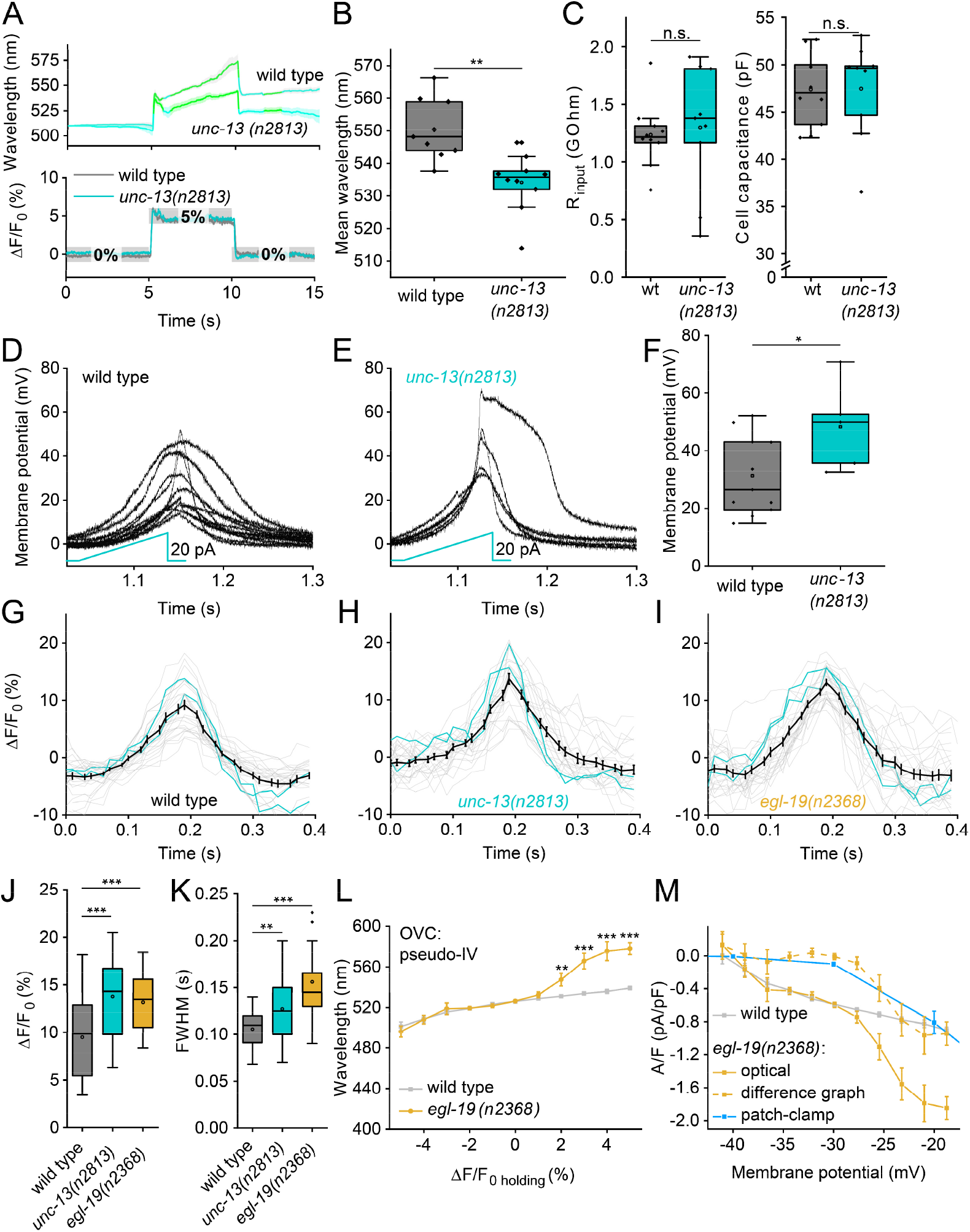
Assessing altered cell physiology in mutants affecting synaptic transmission and ion channels, using the OVC. **(A)** *unc-13(n2813)* mutant and wild type animals were subjected to OVC depolarization, to +5 % ΔF/F_0_. Mean wavelength traces (± S.E.M.), n=9-10. **(B)** Mean wavelength required during the depolarization step to hold +5 % ΔF/F_0_ (p=0.0012). **(C)** Membrane resistance was analyzed in wild type and *unc-13* mutant muscles. n=9-10. **(D, E)** Current ramps (0-20 pA over 100 ms; cyan) were injected into wild type and *unc-13* mutant muscles and the induced voltage increases were aligned to the peak (p=0.74). n=5-11. **(F)** Group data of induced voltage increases in (D and E) (p=0.04). **(G-I)** Optical recordings of spontaneous APs in wild type animals, *unc-13(n2813)* and *egl-19(n2368)* mutants. Overlay of single (each two are highlighted in cyan) and mean (± S.E.M.) traces. n=10-12; 26-28 APs analyzed. **(J, K)** Statistical analysis of peak AP amplitude (wt / *unc-13*: p=8.27E-4, wt / *egl-19* p=3.04E-4) and AP duration (at FWHM; wt / *unc-13*: p=0.0082, wt / *egl-19* p=1.53E-6) of data in (G-I). **(L)** All-optical I/V-relationship obtained in wild type and *egl-19(n2368)* mutants (n=14-17, 2% p=0.018, 3% p=9.36E-9, 4% p=7.26E-10, 5% p=7.92E-9). **(M)** Data in (L) were transformed to I/V-relations, using calibrations in **Fig. 2J, L**. The currents were normalized using capacitance measured in patch-clamped muscle cells (analogous to **Fig. 1I**). Data obtained for wild type (grey) was deduced from *egl-19* data (yellow), generating a difference curve (dashed, yellow), and compared to difference data obtained by electrophysiology^44^ in wild type and *egl-19(n2368)* mutants (blue). Two-sided t-test with Bonferroni correction in (B, C, F, J, K). ***P ≤ 0.001, **P ≤ 0.01, *P ≤ 0.05 and box plots (median, 25-75^th^ quartiles); open dot: mean; whiskers: 1.5x IQR.

*egl-19* g.o.f. mutants were shown to mediate larger currents and slowed inactivation^20,44^. We analyzed APs in the *n2368* allele using voltage imaging (**Fig. 3I, J**), which revealed larger APs of longer duration than in wild type. We compared the two genotypes by generating an optical I/V-curve. In the ± 5 % range (i.e. ca. -40 to -18 mV; **Fig. 2I, J**), *egl-19* mutants and wild type animals were similar below 0 % ΔF/F_0_ (∼resting potential) but diverged significantly once positive ΔF/F_0_ clamp values were reached (i.e. -28 *vs*. -20 mV; **Fig. 3L; Extended data Fig. 8E**). Since we cannot block K^+^-channels in intact worms, we calculated the difference of wild type and mutant data to extract the additional *n2368*-mediated Ca^2+^-currents (**Fig. 3M**). Our optically derived data, corrected for mean membrane capacitance, compared well to electrophysiological data^44^ (again, difference of wild type and *egl-19* mutants; **Fig. 3M**). Thus, EGL-19 g.o.f. channels open at less depolarized membrane potential, compared to wild type EGL-19.

### Optogenetic current-clamp and live-OVC

Our approach enables to achieve an “optogenetic current-clamp”. We wrote according software that allows to present continuous or pulsed wavelength ramps, or a single light pulse of the selected wavelength, while ΔF/F_0_ is recorded live (**Extended data Fig. 9A**). This way, similar to earlier unidirectional approaches^11,12^, we could induce and record APs (or inhibitory potentials) in BWMs in all-optical experiments (**Extended data Fig 9B, C**). To further extend the applicability of the OVC, we wanted to enable the researcher to respond to observations and to dynamically adapt clamping parameters. We thus wrote software “on-the-run” allowing to select holding ΔF/F_0_ values during a running acquisition (**Extended data Fig. 9D**). The software provides a live status update, whether the OVC system is on hold, adapting or if it has reached its limits (**Extended data Fig. 9E, F, Extended data video 2**). Using this tool, we could show that the OVC remained responsive to frequently changing live selected clamping values up to several minutes (**Extended data Fig. 9G, H**).

### Establishing the OVC in cholinergic and GABAergic motor neurons

We tested if the OVC works also in *C. elegans* neurons. Cholinergic and GABAergic motor neurons are small (ca. 2-3 μm cell body, BWMs ca. 50 μm) and exhibit low absolute fluorescence. Both mCerulean (BiPOLES) and QuasAr2 fluorescence were observed in respective ganglia, including the anterior nerve ring and ventral nerve cord (**Fig. 4A, B**). Calibration parameters could be adopted from muscle experiments (**Fig. 4C**). The OVC could readily clamp neuronal voltage-dependent fluorescence between -5 and 5 % ΔF/F_0_ (**Fig. 4C-G**), and also adaptive experiments were possible (**Extended data Fig. 9I, J**).

**Fig. 4.**
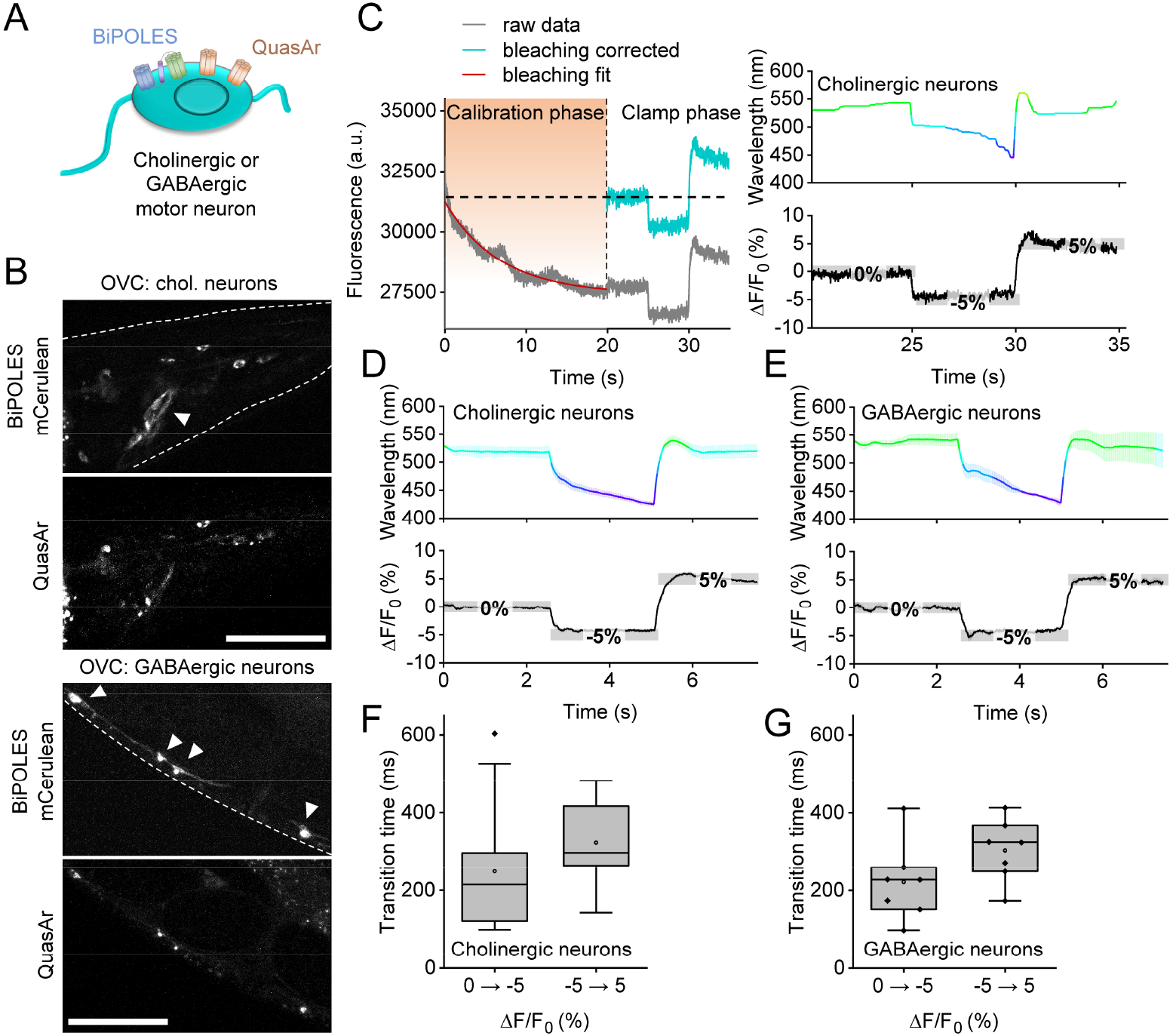
Bi-directional clamping of voltage-dependent fluorescence in *C. elegans* neurons. **(A)** OVC in cholinergic or GABAergic neurons. **(B)** Confocal fluorescence z-projections of BiPOLES (mCerulean) and QuasAr2 expression. Arrowheads: Neuronal cell bodies. Scale bars: 50 μm. **(C)** OVC protocol (holding values 0, -5, 5 % ΔF/F_0_) in cholinergic neurons, single recording. **(D, E)** Mean (± S.E.M.) traces of the OVC in cholinergic (D; n = 14), or GABAergic (E, n = 7) neurons, respectively, holding values: 0, -5, 5 % ΔF/F_0_. **(F, G)** Transition times for 5 and 10 % ΔF/F_0_ steps (F: cholinergic, G: GABAergic neurons); box plots (median, 25-75^th^ quartiles), open dot: mean, whiskers: 1.5x IQR.

### Establishing the OVC in mammalian neurons

Rodent neurons are larger, more hyperpolarized cells and display faster AP kinetics than *C. elegans* muscles and neurons. Since such cells are relevant to studies of human-like neurophysiology, we explored the utility of the OVC in rat hippocampal pyramidal neurons (organotypic slice culture; **Fig. 5A**), expressing QuasAr2 and soma-targeted (som)BiPOLES. After bleaching calibration (**Fig. 5B**), optical clamping could be achieved between ± 3 % ΔF/F_0_ (**Fig. 5B, C**) with the OVC protocol as used in *C. elegans*. Importantly, in the absence of somBiPOLES, monochromator light did not lead to modulation of QuasAr2 fluorescence (**Fig. 5D**), while electrically evoked potential shifts (100 mV depolarization step, from -74.5 mV to +25.5 mV holding voltage) caused clear increases of QuasAr2 fluorescence (ca. 21 % ΔF/F_0_; **Fig. 5E**). In cells expressing QuasAr2 and somBiPOLES, the ± 3 % ΔF/F_0_ OVC protocol caused hyperpolarizing and depolarizing potential jumps (ca. - 4 mV and + 3 mV, respectively; **Fig. 5F, G**). These were small, compared to earlier experiments using somBiPOLES, where 595 nm light application caused depolarization of up to 30 mV, while 400 nm light clamped cells to the Cl^-^ reversal potential^27^. Possibly, due to the more negative resting potential in mammalian neurons compared to *C. elegans* cells, 637 nm activation of Chrimson causes stronger effects, and thus the OVC triggers more compensatory GtACR2 currents. When we expressed somBiPOLES only, 637 nm laser light depolarized hippocampal neurons (via Chrimson) by ca. 21.5 mV, which could be partially counteracted by GtACR2 activation using 530 nm compensation light (**Fig. 5H**). However, due to the high conductance of GtACR2, shunting effects likely prevented further optical hyper- or depolarization. Thus, in mammalian neurons, the OVC works with a limited range, likely due to different voltage and ion conditions in the resting state, and probably due to higher relative expression levels, as compared to *C. elegans*.

**Fig. 5.**
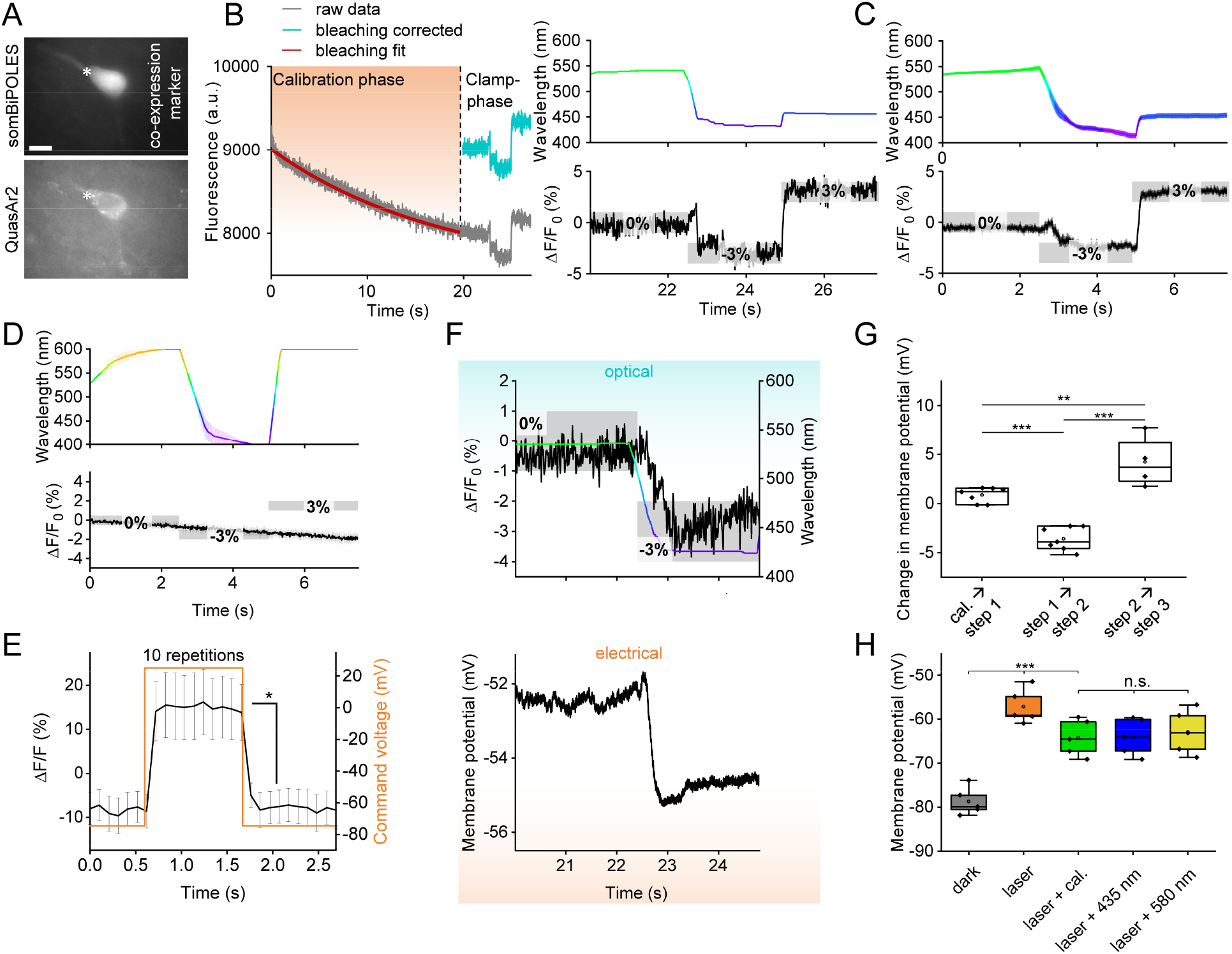
Establishing the OVC in rat hippocampal pyramidal neurons. **(A)** Fluorescence micrographs of somBiPOLES (CFP co-expression marker) and QuasAr2 expression in a neuron (asterisk). Scale bar: 10 μm. **(B)** OVC three-step protocol (0, -3 and 3 % ΔF/F_0_). Right: Wavelength shown in the respective color (upper), holding values and tolerance range (grey boxes, lower panel) are indicated for each step. Left: Orange shade marks transition/calibration period to reach tolerance range. **(C)** Overlay of mean (± S.E.M.) wavelength (upper panel) and fluorescence traces (lower panel; n=12; holding values: 0, -3, 3 % ΔF/F_0_). **(D)** Average fluorescence data (lower panel) for cells expressing QuasAr2 only, while the OVC attempts to run a 0, -3, 3 % ΔF/F_0_ protocol (n = 5; upper panel: Monochromator wavelength). **(E)** QuasAr2 fluorescence during electrically evoked 100 mV depolarization step, from -74.5 mV to +25.5 mV holding voltage. **(F)** Simultaneous patch-clamp (voltage, lower panel) and fluorescence recording (upper panel) with indicated wavelength adaptation and tolerance ranges. **(G)** Statistical analysis of voltage modulation between transition events during simultaneous patch clamp / OVC measurements (n = 3-7). **(H)** Modulation of membrane voltage in cells expressing somBiPOLES only, in response to different light application, as indicated. One-way ANOVA with Bonferroni correction. ***P ≤ 0.001, **P ≤ 0.01, *P ≤ 0.05; n = 3-7. In (G, H), box plots (median, 25-75^th^ quartiles); open dot: mean; whiskers: 1.5x IQR.

### Dynamic suppression of APs in pharyngeal muscle

Thus far, we imposed fixed voltage steps to cells. To see if the OVC can dynamically counteract spontaneous activity, as opposed to shunting hyperpolarization, we turned to a periodically active muscular pump, the *C. elegans* pharynx (**Fig. 6A**).

**Fig. 6.**
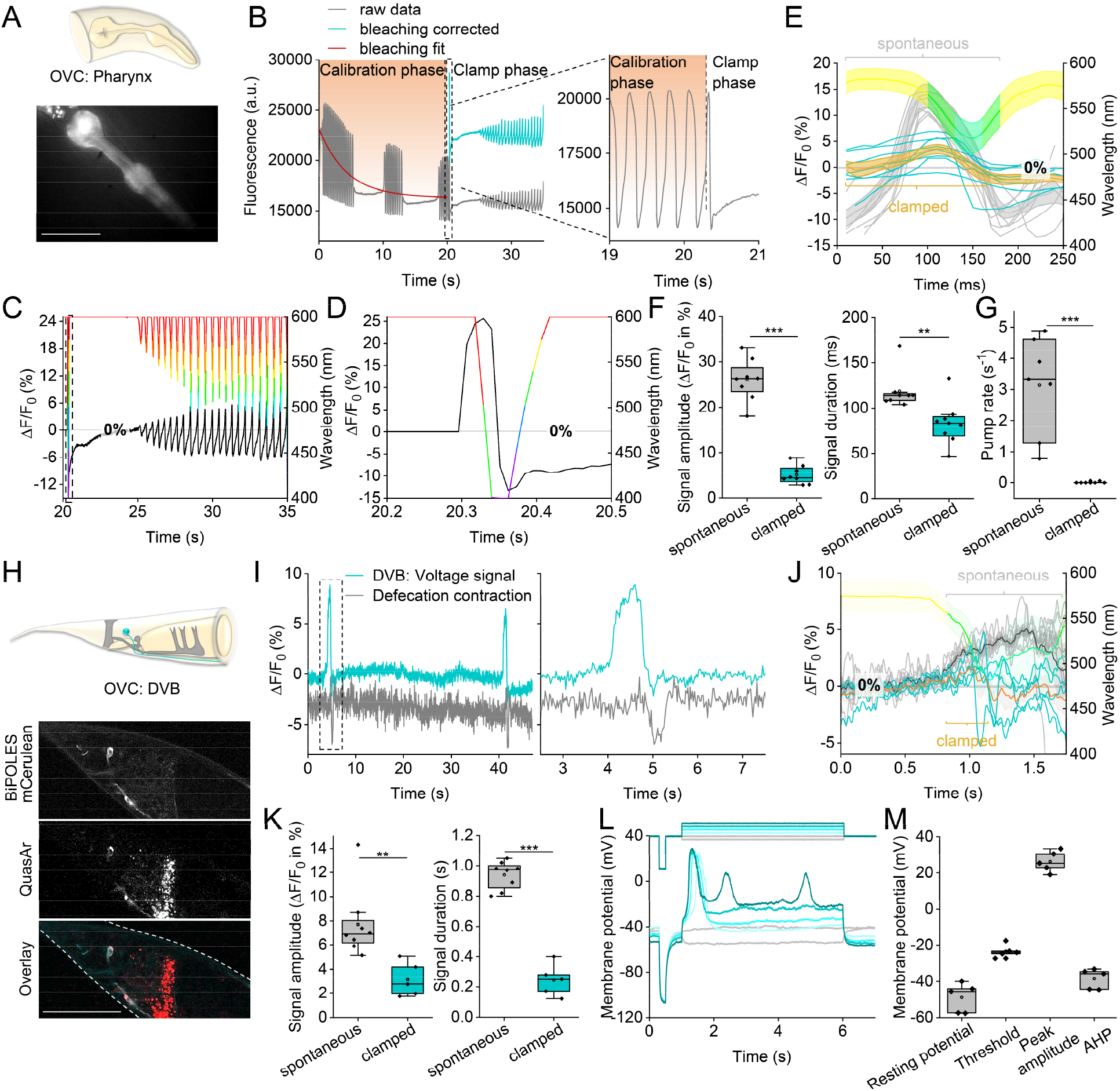
All-optical clamping of APs in pharyngeal muscle and the enteric motor neuron DVB. **(A)** The *C. elegans* pharynx, a muscular pump used for feeding, expressing BiPOLES and QuasAr2 (fluorescence). **(B-D)** OVC experiment in pharyngeal muscle, to hold fluorescence at 0% ΔF/F_0_ and suppress APs. (B) Original trace, calibration and clamping phase. Inset shows close-up of boxed region, transition from calibration to clamp phase. (C) Overlay of wavelength and ΔF/F_0_ traces during clamp phase. (D) Close-up of region boxed in (C): OVC counteracting first AP during clamp phase. (**E**)Aligned traces of spontaneous (light grey, n=8 animals, 5 APs each) and clamped (blue) pharyngeal fluorescence signals. Mean wavelength chosen by the system is shown in the respective color. **(F)** Statistical analysis of data in (E) and additional APs (n=8 animals, 10 APs each; amplitude p = 1.45E-8, duration p = 0.0078), fluorescence voltage signal amplitude and duration at FWHM. **(G)** Pumping, observed visually, was effectively suppressed by dynamic OVC clamping (n=7; p = 2.028E-4). **(H)** The GABAergic motor neuron DVB (cyan) innervates enteric muscles (grey, upper panel). Below: Confocal z-projection of BiPOLES and QuasAr2 fluorescence in DVB, and overlay; scale bar: 50 μm. **(I)** Voltage fluorescence signals of spontaneous APs in DVB (blue trace). Expulsion muscle contraction, deduced from DVB cell body movement, shown as gray trace. Right panel: Close-up of boxed region. **(J)** Overlay of spontaneous (light gray traces, n=8; mean: black) and clamped (blue traces, n=5; mean: orange) DVB voltage signals. Mean wavelength set by the OVC shown in the respective color. **(K)** Statistical analysis of data in (J), mean voltage fluorescence signal amplitude and total duration (amplitude p = 0.0073, duration p = 7.97E-8). **(L)** Current clamp recordings (n=5, lower panel) of DVB in dissected animals, following indicated (upper panel) hyperpolarizing (grey) and depolarizing (cyan) injection of current in 1 pA steps. **(M)** Group data of (L): resting potential, threshold, peak amplitude, duration at FWHM, and afterhyperpolarization (AHP). In (F, G, K and M): box plots (median, 25-75th quartiles), open dot: mean, whiskers: 1.5x IQR. In (F, G, and M), two-sided t-test with Bonferroni correction. ***P ≤ 0.001, **P ≤ 0.01.

This feeding organ exhibits periodic, ca. 4 Hz APs^20^. Closing of the grinder, a structure used to crush bacteria, was forced by activation of *Gt*ACR2 (400 nm), and could be observed by QuasAr2 fluorescence (**Extended data Fig. 10A**). Illumination with only the 637 nm laser caused grinder opening. The OVC could clamp the pharynx statically between -5 and 5 % ΔF/F_0_ (**Extended data Fig. 10B-D**), and dynamically follow its activity to keep it at 0 % ΔF/F_0_ (**Fig. 6B, C; Extended data video 3**). Optically observed APs showed a rise time constant of around 15 ms and duration of ca. 150 ms (at full-width-half-maximum - FWHM)^20^. To clamp these APs dynamically, i.e., to counteract de- and repolarization phases quickly enough, we set the OVC software parameters to respond with more pronounced wavelength changes to a given fluorescence change (increased integral gain; methods, **equation 6**), and reduced the tolerance range to ± 0.005 %. During calibration, APs showed 26.0 ± 1.6 % ΔF/F_0_ mean amplitude and 118.7 ± 7.3 ms duration at FWHM (**Fig. 6B, C, E, F**). Upon clamping, all subsequent attempts to spike were dynamically counteracted and almost completely suppressed (**Fig. 6B-F**). Clamping was rapid: APs occurring at clamping onset were shortened by 65 %, while the OVC traversed the full wavelength spectrum within 20 ms (**Fig. 6D**). Signal amplitude was significantly reduced to ca. 5 ± 0.7 % (**Fig. 6E, F**), as was its duration (to 83.14 ± 8.8 ms at FWHM). The greatly reduced voltage signals also resulted in the suppression of pump events (**Fig. 6G**). This finding supports the effectiveness of OVC-mediated AP suppression. As shown above in BWMs, APs, associated with the opening of the grinder, could be elicited using the current clamp mode (**Extended data Fig. 10E, F**).

### Dynamic clamping APs in the GABAergic motor neuron DVB

*C. elegans* exhibits a rhythmic motor program to move and expel gut contents^29-32^. Expulsion muscle contraction is regulated by an intestinal pacemaker and by GABAergic motor neurons DVB and AVL (**Fig. 6H**). Ca^2+^ imaging of DVB^45^ revealed activity reminiscent of APs, thus far observed in only few *C. elegans* neurons^46^. DVB voltage imaging showed APs (7.7 ± 1 % ΔF/F_0_, 500 ± 60 ms at FWHM) at regular time intervals of 40-50 s, that were followed by posterior body contraction (**Fig. 6I**). Using parameters as for the pharynx, DVB APs were clamped to significantly reduced amplitude and duration (**Fig. 6J, K**; see **Extended data Fig. 10G** for single trace). DVB allowed calibrating the neuronal OVC by patch-clamp electrophysiology^47^: Resting potential was -49.0 ± 8.9 mV (**Fig. 6L, M**), and increasing current steps (1 pA) evoked APs (−23.4 mV threshold, depolarization to 26.3 mV, 359 ms duration (FWHM), -38.5 mV afterhyperpolarization; n=5; **Fig. 6M**). DVB AP amplitude thus is ca. 50 mV. Although fluorescence baseline may be altered by the 637 nm laser and Chrimson activation, this did not evoke APs. Thus, the QuasAr2 signal in DVB, i.e., 7.7 % ΔF/F_0_ for an AP, corresponds to ca. 50 mV (calculating from threshold, or up to 75 mV, from resting potential).

## Discussion

Here, we established the first all-optical voltage-clamp approach to date. We demonstrated its performance in various excitable cell types in intact animals (*C. elegans*) and tested it in mammalian hippocampal neurons. In *C. elegans*, the OVC allowed reliable clamping of voltage in two muscular organs and three neuron types *via* QuasAr2 fluorescence read-out and bidirectional optogenetic actuation *via* BiPOLES. The OVC could further detect the altered postsynaptic excitability of *unc-13* mutants, a homeostatic response to reduced presynaptic input, and allowed deducing all-optical I/V-relationships that provided insight into altered functionality of the L-type VGCC EGL-19 in a g.o,f, mutant, matching electrophysiological data. Using the OVC, we did not only control resting membrane potential, but could also dynamically clamp spontaneous rhythmic activity and APs in pharyngeal muscle and in the motor neuron DVB.

The OVC operates at rates up to 100 Hz on a typical PC with moderate computing power. This orchestrates the communication between camera and monochromator, while in parallel running image acquisition and providing live computation of bleaching-corrected ΔF/F_0_ values, fed into a decision tree and Integral-control algorithm. An alternative algorithm with PID controller and Kalman filter showed similar performance. While the OVC at present is slower than standard patch-clamp electrophysiology, it is not necessarily less sensitive or accurate, within the range of experimental variation, and given variations imposed by dissection required for electrophysiology. The OVC outperforms electrophysiology it in terms of non-invasiveness, throughput and ease of application.

Though activation of Chrimson by QuasAr2 excitation light evoked currents, this effect could be counterbalanced in *C. elegans* by the bidirectionality of BiPOLES, using compensating GtACR2 activation. Despite this low level opening of BiPOLES channels, no significant effects on membrane resistance or membrane potential were observed, intrinsic neuronal activity and cellular excitability were normal, muscles fired regular APs, locomotion behavior was unaltered, and also spontaneous activity in pharyngeal muscle or the DVB neuron were unaffected.

The voltage range covered by the OVC differed between muscles and neurons: 10 % ΔF/F_0_ change corresponded to about 22 mV in BWMs, and to about 65 mV in DVB. Differences in membrane potential and AP amplitude depend on the different ion channels and gradients present in the two cell types, and input resistance. The different range of ΔF/F_0_ fluorescence of QuasAr2 in muscles and neurons may also correspond to different levels of QuasAr2 protein in plasma membrane vs. intracellular membranes of the two cell types. Fluorescence from the latter would not contribute to voltage-dependent ΔF/F_0_, while it would increase overall fluorescence at rest.

BiPOLES mediated currents in a range of approx. 190 pA, comparable to common optogenetic tools^33,34,48^, but falling behind the effects of individually expressed ACR2^19^. In terms of accuracy and in line with previous patch-clamp measurements in BWMs, the examined (patched) cells show a normal distribution in resting potential (ca. -23 to -27 mV), which remains unchanged during the calibration phase. Since our system is based on relative changes in fluorescence, which originate from slightly varying resting potentials, set clamp values may be subject to small variations. The OVC is not a one-to-one replacement for electrophysiology: Membrane potential is mapped as a relative change using fluorescence, thus requiring calibration measurements to deduce absolute values. The rhodopsins used are not as persistent as a patch-clamp electrode and limited by the cells’ reversal potentials for the respective ions. However, in the range of physiological voltage fluctuations in the respective cell type, the OVC is robust and enables generating data with the advantages of a purely optical, contact-less system. This enables acquiring I/V-relationships, and to some extent (since without feedback), current clamp.

In mammalian neurons, optical voltage control was currently limited to a few mV. This was likely due to the resting potential being close to the Cl^-^-reversal potential, but possibly also due to BiPOLES expression levels, as imaging light affected membrane conductance. Thus, in comparison to *C. elegans*, more activation of GtACR2 is required to counteract Chrimson-based depolarization, causing membrane leak. Minimizing the optical crosstalk of voltage sensor and BiPOLES is especially important in mammalian neurons. Further red-shifted voltage sensors and blue-shifted BiPOLES variants are thus required for future OVC experiments. Light-driven ion pumps, provided they become as powerful as the channels used, could be a future alternative.

The speed of the OVC is currently limited by the sensitivity, fluorescence yield, dynamic range of the GEVI (requiring 10 ms exposure times) and its time constant, the camera frame rate, and the computer hardware. With higher fluorescence change in response to membrane potential and a larger absolute GEVI fluorescence (i.e., improved signal to noise ratio), higher frame rates might be achieved. The software currently runs within μManager on a PC. Running the software on an integrated circuit, like a field programmable gate array, and using small ROIs, may enable faster operation speeds up to 1 kHz. GEVI kinetics is unlikely to become a limiting factor, as QuasAr2 can faithfully report on single APs in mammalian neurons (2 ms), clearly showing AP rise and repolarization phases^11^. OFF-kinetics of the actuators may pose a limit to the maximal response times of the OVC: These are ca. 21-45 ms for Chrimson (however, a 10x faster variant has been isolated)^25,26^, and ca. 40 ms for *Gt*ACR2 (here, 20x faster ZipACR was described)^49^. Yet, due to BiPOLES, fast OFF kinetics are less important, as it enables active counteraction. Since channel ON-kinetics are generally faster, they are unlikely to limit the maximal speed achievable by the OVC. That rhodopsin channels show some inactivation during prolonged illumination is not a concern, because the OVC feedback loop can counteract progressive inactivation of BiPOLEs components, at least until one of those would desensitize completely. We observed no problems in measurements lasting up to three minutes.

The OVC detected altered muscular excitability in *C. elegans unc-13* mutants, likely caused by changes in ion channel physiology, and it could directly confirm altered channel excitability and currents through a g.o.f. variant of the EGL-19 VGCC. This shows that the OVC represents an approach to enable all-optical, contact-less high-throughput screening applications, e.g. of compound libraries targeted at ion channels. Testing its use in mammalian cell types other than neurons, where such ion channels can be expressed individually, and where the resting membrane potential is less depending on the Cl^-^-gradient, will facilitate this approach.

The establishment of the OVC paves the way for all-optical control of individual neurons in freely behaving transparent animals like *C. elegans*. Accuracy of fluorescence quantification (and thus OVC feedback) requires the cell to remain in focus, which can be achieved with the OVC, while it is impossible to keep a patch-clamp electrode physically attached to such a small animal. Using the OVC on neurons controlling behavior will allow fine-tuning of behavioral aspects, and enable understanding how activity of the individual cell regulates them. Online behavioral analysis (e.g. extent of body bending, locomotion velocity), may be used for feedback to the neuron such that behavior can be dynamically controlled.

Transferring the OVC to other model organisms may require modifications depending on the respective cellular properties. Extending electrophysiology applications, and provided the monochromator light is projected via a digital micromirror device, the OVC should enable efficient space clamp, as well as dynamic local clamping in neuronal processes. The OVC software is universally applicable, as it can be adapted to other GEVI-optogenetic actuator combinations. The OVC broadens applicability of optogenetics as it allows modulation in closed-loop, to better adapt to the variable activity patterns found in living organisms. Dynamic responsiveness is also advantageous with regard to future therapeutic applications, for example in acute control of seizures^50^, or in adaptive deep brain stimulation, as it would allow adjusting the therapy to the need of the patient^51^.

## Acknowledgements

We thank Jiajie Shao for support in confocal microscopy, Adam Cohen for reagents, and Sven Plath, Regina Wagner, Hans-Werner Müller, Franziska Baumbach, Nico Sturman, Adrian Breicher, Timotheus Kozlowski, Rolf Bergs, and Kathrin Sauter for expert technical assistance.

## Funding

Goethe University (AG); Deutsche Forschungsgemeinschaft, SFB807/B02 (AG), SPP1926-XIb / WI 4485/3-2 (JSW), and SFB1315/C01 (JV, PH); Max Planck Society, PhD fellowship (ACFB); Chan Zuckerberg Initiative, support (QL, CIB)

## Author contributions

Conceptualization: ACFB, AG, JSW

Methodology: ACFB, AB, JV, PH, JSW

Investigation: ACFB, JFL, SRR, QL, AN, CW, NZ, HD, MJ

Visualization: ACFB, AG

Funding Acquisition: AG, ACFB, JSW, PH, CIB

Project administration: AG

Supervision: AG, JSW, CIB

Writing – original draft: ACFB, AG

Writing – review & editing: ACFB, AG, SRR, JSW, JV, CIB, PH

## Competing Interests

A patent application has been filed: EP 21162331.9 (ACFB, AG).

## Data and materials availability

The datasets generated during and/or analyzed during the current study are available from the corresponding author on reasonable request. This also applies to materials described in the study.

## Software availability

The software / code used for operating the OVC is written in beanshell, the scripting language used by μManager freeware. Scripts (“OVC_main_script”, “OVC_on_the_run_script” “OVC_4_step_script”, “OVC_pseudo-IV_script”, “optical_current_clamp_script”) are available as a supplementary files. They will be made available upon publication under the license XYZ.

## Extended data

Online Methods

Extended data Figures 1 to 10

Extended data videos 1 to 3

## Online Methods

### Transgenic *C. elegans* strains

The following strains were used or generated, and are available upon request: **ZX2476**: *zxEx1139[pmyo-3::QuasAr2; pmyo-2::CFP]*, **ZX2482**: *zxEx1145[pmyo-3::QuasAr2; pmyo-2::CFP]; zxIs5[punc-17::ChR2(H134R)::yfp;lin-15+]*, **ZX2483**: *zxEx1146[punc-17::ACR2::eYFP; pmyo-3::QuasAr2; pelt-2::GFP]*, **ZX2586**: *zxEx1228[punc-17::GtACR2::mCerulean::*β*HK::Chrimson; pelt-2::GFP]*, **ZX2714**: *zxEx1250[punc-17::GtACR2::mCerulean::*β*HK::Chrimson; pmyo-3::QuasAr2; pelt-2::GFP]*, **ZX2753**: *zxEx1266[pmyo-3::GtACR2::mCerulean::*β*HK::Chrimson; pmyo-3::QuasAr2; pmyo-2::CFP]*, **ZX2755**: *zxEx1268[punc-47::QuasAr2::GFP; pmyo-2::CFP]*, **ZX2826**: *zxEx1282[pmyo-2::QuasAr2; pmyo-2::GtACR2::mCerulean::*β*HK::Chrimson; pmyo-3::CFP]*, **ZX2827**: *zxEx1283[punc-17::GtACR2::mCerulean::*β*HK::Chrimson; punc-17::QuasAr2; pelt-2::GFP]*, **ZX2828**: *zxEx1284[punc-47::QuasAr2::GFP; punc-47::GtACR2::mCerulean::*β*HK:: Chrimson; pmyo-2::CFP]*, **ZX2876**: *zxIs139[pmyo-3::GtACR2::mCerulean::*β*HK::Chrimson; pmyo-3::QuasAr2; pmyo-2::CFP]*, **ZX2935**: *unc-13(n2813)*; *zxIs139[pmyo-3::GtACR2::mCerulean::*β*HK::Chrimson; pmyo-3::QuasAr2; pmyo-2::CFP]*.

### Molecular biology

Plasmids pAB4 (p*unc-17*::ACR2::eYFP), pAB16 (p*myo-3*::QuasAr; Addgene plasmid #130272), pAB17 (p*unc-17*::QuasAr), pAB23 (p*tdc-1s*::QuasAr::GFP) and pNH12 (p*myo-2*::MacQ::mCitrine) were described earlier ^19, 20^. **pAB26 (p*unc-17*::GtACR2::mCerulean::βHK::Chrimson)** was generated by Gibson Assembly based on RM#348p (p*unc-17*; a gift from J. Rand) and pAAV-hSyn-BiPOLES-mCerulean (Addgene plasmid #154944), using *Nhe*I and primers 5’-attttcaggaggacccttggATGGCATCACAGGTCGTC-3’ and 5’-ataccatggtaccgtcgacgTCACACTGTGTCCTCGTC-3’. **pAB27 (p*myo-3*::GtACR2::mCerulean::βHK::Chrimson)** was generated *via* Gibson Assembly based on pDD96.52 (p*myo-3*, Addgene plasmid #1608) and pAAV-hSyn-BiPOLES-mCerulean, using *Bam*HI and primers 5’-actagatccatctagagATGGCATCACAGGTCGTC-3’ and 5’-ttggccaatcccgggCACTGTGTCCTCGTCCTC-3’. **pAB28 (p*unc-47*::QuasAr::GFP)** was generated by Gibson Assembly based on pMSM08 (p*unc-47*::eGFP::MmBoNTB) and pAB23 (p*tdc-1s*::QuasAr::GFP), using *Xma*I, *Msc*I and primers 5’-ttacagcaccggtggattggATGGTAAGTATCGCTCTG-3’ and 5’-ttctacgaatgctcctaggcCTATTTGTATAGTTCATCCATGC-3’. **pAB29 (p*myo-2*::QuasAr)** was generated by Gibson Assembly based on pNH12 (p*myo-2*::MacQ::mCitrine) and pAB16 (p*myo-3*::QuasAr), using primers 5’-caccgagtgaGAAGAGCAGGATCACCAG-3’, 5’-tgcagagcgatacttaccatCCCCGAGGGTTAAAATGAAAAG-3’, 5’-ATGGTAAGTATCGCTCTGCAG-3’ and 5’-cctgctcttctcaCTCGGTGTCGCCCAGAATAG-3’. **pAB30 (p*myo-2*::GtACR2::mCerulean::bHK::Chrimson)** was generated by Gibson Assembly based on pNH12 (p*myo-2*::MacQ::mCitrine) and pAAV-hSyn-BiPOLES-mCerulean, using *Bam*HI, *Hind*III and primers 5’-ggacgaggacacagtgtgaaAAGAGCAGGATCACCAGC-3’ and 5’-agacgacctgtgatgccatgCCCCGAGGGTTAAAATGAAAAG-3’. **pAB31 (p*unc-47*::GtACR2::mCerulean::bHK::Chrimson)** was made by Gibson Assembly based on pAB28 (p*unc-47*::QuasAr::GFP) and pAAV-hSyn-BiPOLES-mCerulean, using *Age*I, *Eco*RI and primers 5’-acatttatttcattacagcaATGGCATCACAGGTCGTC-3’ and 5’-agcgaccggcgctcagttggTCACACTGTGTCCTCGTC-3’.

For neuronal expression, the coding sequence of QuasAr2 ^52^ (Addgene #107705) was cloned together with a trafficking signal (ts: KSRITSEGEYIPLDQIDINV) and an ER-export signal (ER: FCYENEV) from the Kir 2.1 channel ^53, 54^ into an AAV2-backbone behind a human synapsin (hSyn) promoter resulting in **pAAV-QuasAr2-ts-ER**.

### OVC voltage imaging experiments

For voltage imaging experiments, animals were supplemented with all-*trans* retinal (ATR; Sigma-Aldrich, USA): One day prior to experiments, transgenic L4 stage animals were transferred to NGM plates, seeded with OP50 bacterial suspension mixed with ATR (stock in ethanol). To avoid interfering fluorescence of unbound ATR, its concentration was adjusted for each tissue. Final ATR concentrations (mM): BWMs (0.01), pharynx (0.03), cholinergic neurons (0.1), GABAergic neurons (0.005). Animals were immobilized with polystyrene beads (0.1 μm diameter, at 2.5 % w/v, Sigma-Aldrich) on 10 % agarose pads (in M9 buffer). Voltage-dependent fluorescence of QuasAr2 was excited with a 637 nm red laser (OBIS FP 637LX, Coherent) at 1.8 W/mm^2^ and imaged at 700 nm (700/75 ET Bandpass filter, integrated in Cy5 filter cube, AHF Analysentechnik), while optogenetic actuators (BiPOLES, *Gt*ACR2 or ChR2(H134R)) were activated using a monochromator (Polychrome V, Till Photonics / Thermo Scientific), set to emit light from 400 to 600 nm at 300 μW/mm^2^. Imaging was performed on an inverted microscope (Zeiss Axio Observer Z1), equipped with a 40x oil immersion objective (Zeiss EC Plan-NEOFLUAR 40x/ N.A. 1.3, Oil DIC ∞ / 0.17), a laser beam splitter (HC BS R594 lambda/2 PV flat, AHF Analysentechnik), a galilean beam expander (BE02-05-A, Thorlabs) and an EMCCD or an sCMOS camera (Evolve 512 Delta, Photometrics, or Kinetix 22, Teledyne). All OVC experiments were performed at up to 100 fps with 10 ms exposure and a binning of 4×4 (computer specifications: 24 GB RAM, AMD FX-8150 Octa-core processor (3.6 GHz), NVIDIA GeForce GT 520). To induce pharyngeal pumping, animals were incubated in 3 μl serotonin (20 mM, in M9) for 3 min prior to experiments.

### OVC software

The OVC control software was written in Beanshell, the scripting language used by the open source microscopy software μManager v. 1.4.22 ^35^. Scripts are provided in supplementary information. Experiments were initiated via the μManager script panel. For support of Polychrome V related commands, a copy of the Java archive file *TILLPolychrome*.*jar* must be placed into the MMPlugins folder and the dynamic link library file *TILLPolychromeJ*.*dll* into the Sys32 folder (both provided by Till Photonics). The main OVC software is compatible with all cameras supported by μManager. Before running the software, simple rectangular ROIs (to save computation capacity, **Extended data Fig. 1B**) must be selected in the live image mode by using the ROI button in the μManager main window. Once a ROI is selected, the script can be executed via the script panel GUI. An input tab prompts the user to select the OVC- and acquisition parameters. The software allows to define a holding ΔF/F_0_ value (in %) and the number of frames for each of the steps (three- or four-step-protocol). One can further select the number of frames for the calibration period, the tolerance range (in %), the algebraic sign of the increment factor that decides whether to increase or decrease the wavelength with respect to the current ΔF/F_0_, the starting wavelength and the wavelength limits (in nm). The free choice of these parameters ensures that the program can also be used for other and/or future combinations of optogenetic tools with different spectral properties. For acquisition, light intensity of the Polychrome V (in %), exposure time of the camera (in ms) and binning can be selected.

During calibration, acquired grey values are stored in an array and used to evaluate the parameters for the exponential decay function to correct for bleaching of the voltage sensor, live during the clamping phase (**Equation 1-3**; based on the ImageJ Plugin “Correctbleach” ^55, 56^). At the last time point of the calibration phase, the first 50 bleaching corrected gray values of the recording are used to calculate F_0_. Subsequently, bleaching corrected ΔF/F_0_ values are calculated at each time point of the clamping phase (**Equation 4**). Once the system has access to the bleaching-corrected ΔF/F_0_ values, it feeds them into a decision tree algorithm, where they are compared to a desired holding ΔF/F_0_ value. Depending on the sign of the difference, it decides, whether to shift the wavelength blue or red (mirrored by the sign of the increment i[k]). Wavelength adaptation occurs *via* an implemented I-controller (integral gain K_I_ was empirically chosen to fit the desired system-behavior) that maps the “I” dynamics by buffering the manipulated variable (wavelength *λ*) to compensate for permanent control deviation, which would otherwise occur by using a sole P-controller (**Extended data Fig. 1A**). That means that the difference (error) between command and current ΔF/F_0_ determines the wavelength change of the monochromator – hence, the current wavelength results from the previous plus the respective wavelength change at each time point (**Equation 5 and 6**). The software comes with a three- or four-step protocol, where among others, the desired holding values (within a tolerance range) and number of frames for each step can be selected by the user (**Extended data Fig. 1C**). The system changes the wavelength only if the tolerance range is exceeded.

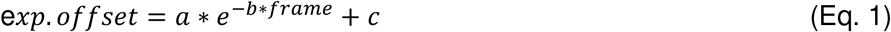

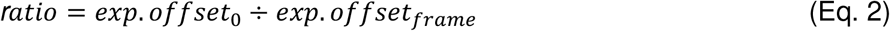

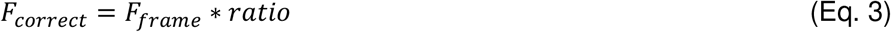

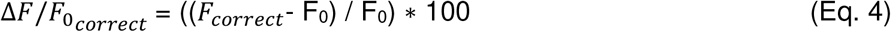

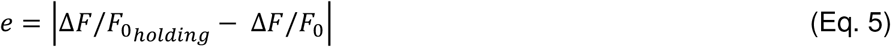

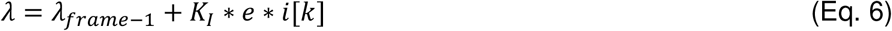

Exp. offset: exponential offset. a, b, c: exponential fitting parameters. Ratio: ratio between first and current exponential offset. *F*_*correct*_: Bleaching-corrected gray values. *F*_*frame*_: Gray value of current frame. 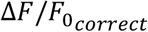: Bleaching-corrected relative change in fluorescence. F_0_: Resting fluorescence value. *e*: error. *λ*: wavelength. *K*_*I*_: integral gain. i[k]: current sign of the control error. Equations 1-3 based on the ImageJ Plugin “Correctbleach”^55^.

When the OVC acquisition is complete, a results text file is given out, and the respective image stack is saved automatically as TIFF file. The results file comprises the selected parameters, the bleach correction function and its quality, the framerate, as well as time, (bleaching corrected-) raw grey- and ΔF/F_0_ values, and wavelength traces. In addition, status information (system on hold, adapting, or wavelength limits reached) is recorded for each time point of the clamp phase. For the “on-the-run” mode, an additional control window opens as soon as the measurement starts. Here, the holding ΔF/F_0_ values can be selected live, either via a range slider or with help of the keyboard arrow keys, while a live status update is displayed.

In addition, “pseudo I/V curve” software is available to clamp relative fluorescence in 11 consecutive steps based on previously selected upper and lower limits, resulting in an output of the average ΔF/F_0_ value and associated wavelength achieved for each step (calculated as a mean of the last 25 % of each step). An optical wavelength / ΔF/F_0_ curve is produced and can be translated into an estimated I/V-diagram using the following linear calibration functions (**Fig. 2J, L**):

linear regression: y = a + bx + e, where e is assumed as i.i.d. residuals with mean 0.

Membrane potential as a function of ΔF/F_0_ holding (estimate: all observations, n = 80, w/o outliers)^a^

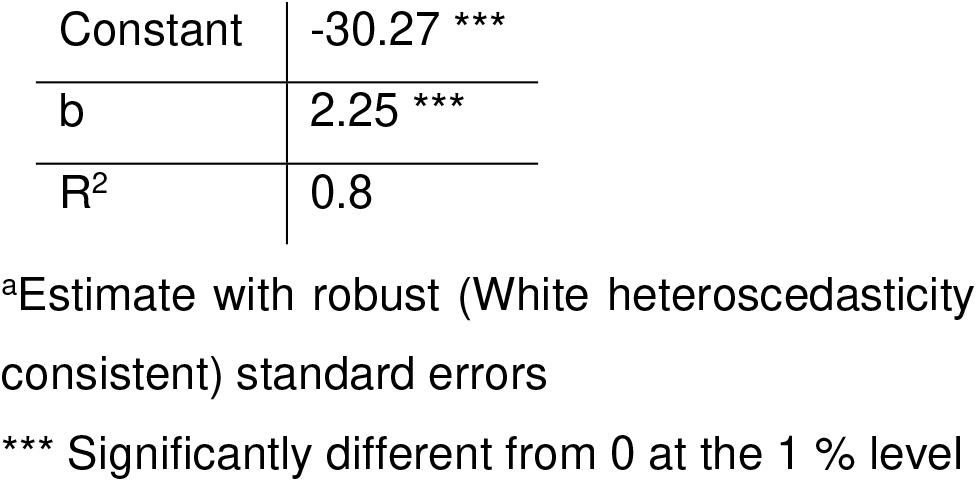

linear regression: y = a + bx + e, where e is assumed as i.i.d. residuals with mean 0

Current as a function of wavelength (estimate: all observations, n = 72)a

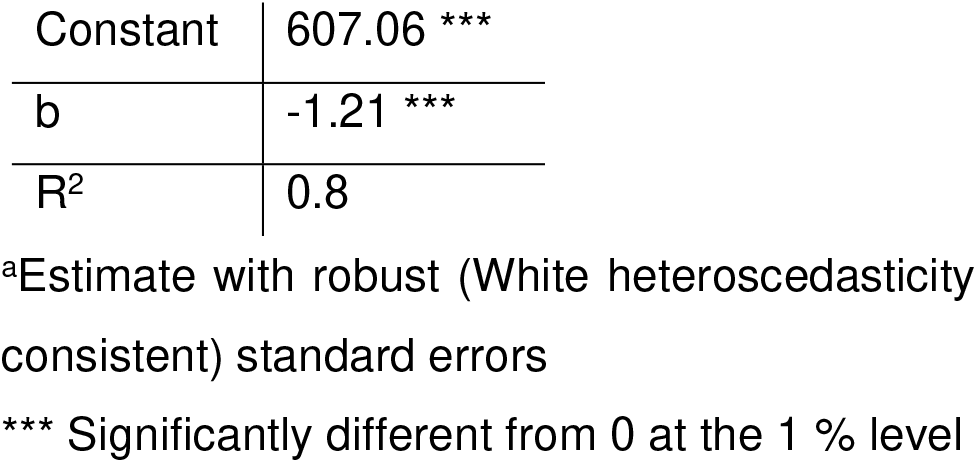

A further software version involves a PID-controller to adjust the wavelength of the OVC. At each time point, the new wavelength is calculated by the addition of the output (u) of the PID equation to the calibration wavelength (**Equations 7-9**).

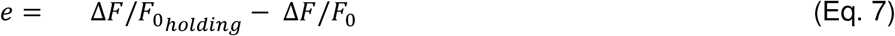

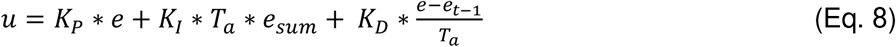

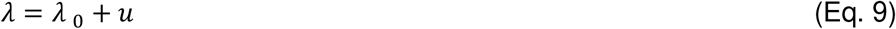

*e*: current error. *u*: output. *K*_*P*_: proportional gain. *K*_*I*_: integral gain. *K*_*D*_: derivative gain. *e*_*t-1*_: previous error. *T*_*a*_: sampling rate. *e*_*sum*_: summed up error. *λ* _0_: calibration wavelength.

Parameter tuning was performed by applying the Ziegler-Nichols method to empirically tune the P,I and D control gains for better control-performance (**Extended data Fig. 2C**). Therefore, *K*_*P*_ was adapted until the system started to oscillate at the critical proportional gain (*K*_*P crit*_). The oscillation period was noted as *T*_*crit*_. PID coefficients were calculated according to **Equations 10 and 11**:

#### Ziegler-Nichols parameter tuning

|  |  |  |
| --- | --- | --- |
| $K_P$ | $T_n$ | $T_v$ |
| $0.6 K_{P\text{crit}}$ | $0.5 T_{\text{crit}}$ | $0.12 T_{\text{crit}}$ |

$K_P$ : proportional gain. $K_{P\text{crit}}$ : critical proportional gain. $T_{\text{crit}}$ : period of oscillation. $T_n$ : integration-coefficient. $T_v$ : derivation-coefficient<sup>36</sup>.

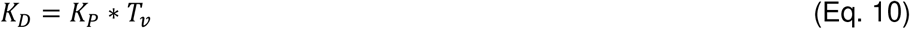

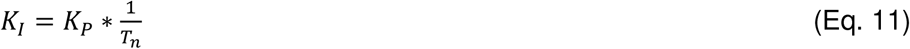

To also achieve a better control performance, a Kalman filter for sensor-signal smoothing was added (**Extended data Fig. 2A**). Here, the error covariances of the measurement- and system noise of the Kalman filter were empirically adjusted so that the OVC controller dynamics achieved a desired performance. After initializing the system state with an a priori estimate (variables 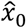: estimated fluorescence at the beginning of the measurement and *P*_0_: estimated error covariance of measured fluorescence vs. estimated fluorescence at the beginning of the measurement), prediction and correction are performed at each time step, alternately propagating the system state in time, and correcting it with new observations (**Equations 12-17**) ^57^:

1. Prediction of the fluorescence value and the error covariance:

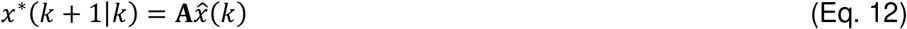

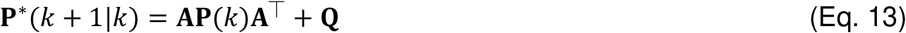

*k*: discrete time-Step. *A*: system matrix, was here set to scalar value one (which is sufficient for error smoothing). *x*^*^: predicted fluorescence. *P*^*^: predicted error covariance of the estimate. *Q*: Error covariance of the system noise. *P(k)*: estimated error covariance.
2. Calculation of the Kalman gain:

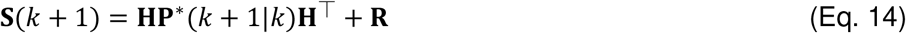

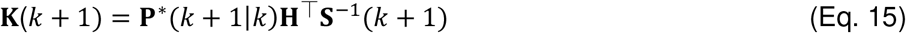

S: auxiliary quantity for determining the Kalman gain. H: system output matrix. R: error covariance matrix of the measurement noise. K: Kalman gain.
3. Correction of the state estimate:

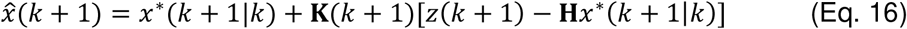

*z(k + 1)*: current measurement. 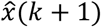: predicted measurement. *x(k + 1)*: current estimate.
4. Correction of the covariance estimation

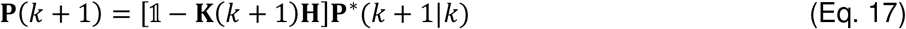

The system matrices A and H were set as constant scalars (A = 1, H = 1; constant system dynamics) since this assumption is perfectly sufficient for signal smoothing for the PID controller. Here, the sampling rate is fast enough that the Kalman filter can respond to membrane potential changes occurring in this time period with reasonable estimates.

Another software, “optical current clamp” is used for the purely optical adaptation of a current clamp. The experimenter has the choice to set single pulses of certain wavelength and duration or to select step-like or continuous wavelength ramps prior to the experiment. Similar to the main OVC script, this software provides bleaching correction and live ΔF/F_0_ readout.

### Quality assessment of the OVC

The predictive accuracy for bleaching correction had R^2^ values above 0.95 in most cases, and at least above 0.8 (**Extended data Fig. 6A**). Experiments with lower R^2^ values were attributable to APs occurring during the calibration phase (e.g. in pharyngeal muscle). As expected, the fraction of system saturation decreased once optogenetic tools for de- and hyperpolarization were simultaneously used, which stresses the importance of a second, opposing actuator (**Extended data Fig. 6B**). While for the single-tool approaches, the percentage of system saturation was relatively high with 16.3 ± 2.8 %, this relation was significantly reduced for all bidirectional BiPOLES combinations. This was also reflected in the speed of the system, where compared to single-tool combinations, BiPOLES was significantly faster, particularly for driving excited or hyperpolarized states back towards resting potential (**Extended data Fig. 6C**).

### Electrophysiology in *C. elegans*

Electrophysiological recordings from body wall muscle cells were conducted in immobilized and dissected adult worms as described previously^41^. Animals were immobilized with Histoacryl L glue (B. Braun Surgical, Spain) and a lateral incision was made to access neuromuscular junctions (NMJs) along the anterior ventral nerve cord. The basement membrane overlying body wall muscles was enzymatically removed by 0.5 mg/ml collagenase for 10 s (C5138, Sigma-Aldrich, Germany). Integrity of BWMs and nerve cord was visually examined *via* DIC microscopy. Recordings from BWMs were acquired in whole-cell patch-clamp mode at 20-22°C using an EPC-10 amplifier equipped with Patchmaster software (HEKA, Germany). Voltage clamp experiments were conducted at the given holding potential (e.g. -24 mV). Membrane potentials in body wall muscle cells were recorded in current clamp mode. In regular recordings no additional current pulse was injected. For measurements of input resistance, a current pulse of -20 pA was injected via the Patchmaster software for 1000 ms in five consecutive pulses with 5 s breaks in between the current pulses.

The head stage was connected to a standard HEKA pipette holder for fire-polished borosilicate pipettes (1B100F-4, Worcester Polytechnic Institute, Worcester, MA, USA) of 4-10 MΩ resistance. The extracellular bath solution consisted of 150 mM NaCl, 5 mM KCl, 5 mM CaCl_2_, 1 mM MgCl_2_, 10 mM glucose, 5 mM sucrose, and 15 mM HEPES (pH 7.3 with NaOH, ≈330 mOsm). The internal/patch pipette solution consisted of K-gluconate 115 mM, KCl 25 mM, CaCl_2_ 0.1 mM, MgCl_2_ 5 mM, BAPTA 1 mM, HEPES 10 mM, Na_2_ATP 5 mM, Na_2_GTP 0.5 mM, cAMP 0.5 mM, and cGMP 0.5 mM (pH 7.2 with KOH, ≈320 mOsm). For some experiments, BAPTA was replaced by EGTA (5 mM).

For light activation a monochromator (Polychrome V, Thermo Scientific) was used, ranging from 400 to 600 nm at 300 μW/mm^2^. To create conditions as similar as possible to those for the OVC experiments, additional excitation was performed with a laser at 637 nm. Subsequent analysis and graphing were performed using Patchmaster, and Origin (Originlabs).

Electrophysiological recordings from the DVB neuron were conducted in immobilized and dissected adult worms as described previously ^46^ with minor modifications. Both dissection and recording were performed at room temperature. Briefly, an animal was immobilized with cyanoacrylate adhesive (Vetbond tissue adhesive; 3M) along the ventral side of the posterior body on a Sylgard 184-coated (Dow Corning) glass coverslip. A small longitudinal incision was made using a diamond dissecting blade (Type M-DL 72029 L; EMS) in the tail region along the glue line. The cuticle flap was folded back and glued to the coverslip with GLUture Topical Adhesive (Abbott Laboratories), exposing DVB. The coverslip with the dissected preparation was then placed into a custom-made open recording chamber (∼1.5 ml volume) and treated with 1 mg/ml collagenase (type IV; Sigma) for ∼10 s by hand pipetting. The recording chamber was subsequently perfused with the standard extracellular solution using a custom-made gravity-feed perfusion system for ∼10 ml. The standard pipette solution was (all concentrations in mM): K-gluconate 115; KCl 15; KOH 10; MgCl_2_ 5; CaCl_2_ 0.1; Na_2_ATP 5; NaGTP 0.5; Na-cGMP 0.5; cAMP 0.5; BAPTA 1; Hepes 10; Sucrose 50, with pH adjusted to 7.2, using KOH, osmolarity 320–330 mOsm. The standard extracellular solution was: NaCl 140; NaOH 5; KCl 5; CaCl_2_ 2; MgCl_2_ 5; Sucrose 15; Hepes 15; Dextrose 25, with pH adjusted to 7.3, using NaOH, osmolarity 330–340 mOsm. Liquid junction potentials were calculated and corrected before recording. Electrodes with resistance of 20–30 MΩ were made from borosilicate glass (BF100-58-10; Sutter Instruments) using a laser pipette puller (P-2000; Sutter Instruments) and fire-polished with a microforge (MF-830; Narishige). Recordings were performed using single-electrode whole-cell current-clamp (Heka, EPC-10 USB) with two-stage capacitive compensation optimized at rest, and series resistance compensated to 50 %.

### Preparation of organotypic hippocampal slice cultures and transgene delivery

All procedures were in agreement with the German national animal care guidelines and approved by the independent Hamburg state authority for animal welfare (Behörde für Justiz und Verbraucherschutz). They were performed in accordance with the guidelines of the German Animal Protection Law and the animal welfare officer of the University Medical Center Hamburg-Eppendorf.

Organotypic hippocampal slices were prepared from Wistar rats of both sexes (Jackson-No. 031628) at post-natal day 5-7 as previously described ^58^. In brief, dissected hippocampi were cut into 350 μm slices with a tissue chopper and placed on a porous membrane (Millicell CM, Millipore). Cultures were maintained at 37°C, 5% CO_2_ in a medium containing 80% MEM (Sigma M7278), 20% heat-inactivated horse serum (Sigma H1138) supplemented with 1 mM L-glutamine, 0.00125% ascorbic acid, 0.01 mg ml^-1^ insulin, 1.44 mM CaCl_2_, 2 mM MgSO_4_ and 13 mM D-glucose. No antibiotics were added to the culture medium. Pre-warmed medium was replaced twice per week.

For transgene delivery, organotypic slice cultures were transduced at DIV 3-5 with a recombinant adeno-associated viral vector (rAAV) encoding a soma-targeted version of BiPOLES ^27^ (AAV9-CaMKII-somBiPOLES-mCerulean, Addgene #154948). The rAAV was locally injected into the CA1 region using a Picospritzer (Parker, Hannafin) by a pressurized air pulse (2 bar, 100 ms) expelling the viral suspension into the slice ^59^. During virus transduction, membranes carrying the slices were kept on pre-warmed HEPES-buffered solution. At DIV 14-16, individual CA1 pyramidal cells in the previously somBiPOLES-transduced slices were transfected by single-cell electroporation ^60^ with the plasmid pAAV-QuasAr2-ts-ER at a final concentration of 10 ng/μl in K-gluconate-based solution consisting of (in mM): 135 K-gluconate, 10 HEPES, 4 Na_2_-ATP, 0.4 Na-GTP, 4 MgCl_2_, 3 ascorbate, 10 Na_2_-phosphocreatine (pH 7.2). An Axoporator 800A (Molecular Devices) was used to deliver 50 hyperpolarizing pulses (−12 V, 0.5 ms) at 50 Hz. During electroporation slices were maintained in pre-warmed (37°C) HEPES-buffered solution (in mM): 145 NaCl, 10 HEPES, 25 D-glucose, 2.5 KCl, 1 MgCl_2_ and 2 CaCl_2_ (pH 7.4, sterile filtered). A plasmid encoding hSyn-mCerulean (at 50 ng/μl) was co-electroporated and served as a marker to rapidly identify cells putatively co-expressing QuasAr2 and somBiPOLES under epifluorescence excitation.

### OVC voltage imaging and electrophysiology experiments in rat organotypic slices

At DIV 19-21, voltage imaging and/or whole-cell patch-clamp recordings of transfected CA1 pyramidal neurons were performed. Experiments were done at room temperature (21-23°C) using a BX51WI upright microscope (Olympus) equipped with an EMCCD camera (Evolve 512 Delta, Photometrics), dodt-gradient contrast and a Double IPA integrated patch amplifier controlled with SutterPatch software (Sutter Instrument). QuasAr2 was excited with a 637 nm red laser (OBIS FP 637LX, Coherent) at 0.52 W/mm^2^ via a dichroic mirror (660nm, Edmund Optics) and voltage-dependent fluorescence was detected through the objective LUMPLFLN 60XW (Olympus) and through an emission filter (732/68 nm Bandpass filter, Edmund Optics). somBiPOLES was activated using a monochromator (Polychrome V, Till Photonics / Thermo Scientific), set to emit light from 400 to 600 nm at 1.7 mW/mm^2^. Laser and monochromator light were combined using a 605nm dichroic mirror (Edmund Optics). Irradiance was measured in the object plane with a PM400 power meter equipped with a S130C photodiode sensor (Thorlabs) and divided by the illuminated field of the Olympus LUMPLFLN 60XW objective (0.134 mm^2^). As for *C*.*elegans*, OVC experiments were performed at ≈ 90 fps with 10 ms exposure and a binning of 4×4.

For electrophysiology, patch pipettes with a tip resistance of 3-4 MOhm were filled with intracellular solution consisting of (in mM): 135 K-gluconate, 4 MgCl_2_, 4 Na_2_-ATP, 0.4 Na-GTP, 10 Na_2_-phosphocreatine, 3 ascorbate, 0.2 EGTA, and 10 HEPES (pH 7.2). Artificial cerebrospinal fluid (ACSF) consisted of (in mM): 135 NaCl, 2.5 KCl, 2 CaCl_2_, 1 MgCl_2_, 10 Na-HEPES, 12.5 D-glucose, 1.25 NaH_2_PO_4_ (pH 7.4). Synaptic transmission was blocked by adding 10 μM CPPene, 10 μM NBQX, and 100 μM picrotoxin (Tocris) to the extracellular solution. Measurements were corrected for a liquid junction potential of -14,5 mV. Access resistance of the recorded neurons was continuously monitored and recordings above 30 MΩ were discarded.

### Statistical methods

Statistical methods used are described in the figure legends. No sample was measured multiple times, and no samples were excluded from analysis. Statistical analysis was performed in Prism (Version 8.0.2, GraphPad Software, Inc.), OriginPro 2021 (OriginLab Corporation, Northampton, USA), Microsoft Excel 2016, 2019, SutterPatch V2 (Sutter Instrument) MATLAB 2016b, 2019b (Mathworks), Stata 12 ic, or ImageJ v1.53c. No statistical methods were applied to predetermine sample size. However, sample sizes reported here are consistent to data presented in previous publications, and which typically allow distinction of significant differences in the types of experiments used. Throughout, n indicates biological replicates (i.e. animals analyzed), with additional mention of specific data that was analyzed (e.g. action potential number), where appropriate. Data was tested for normality prior to statistical inference. Data is given as means ± SEM, if not otherwise stated. Significance between data sets after paired or two-sided t-test or ANOVA, with Bonferroni correction for multiple comparisons, is given as p-value (* p ≤0.05; ** p ≤ 0.01; *** p ≤ 0.001), when not otherwise stated. Group data are presented as box plots with median (line), 25-75th quartiles (box), mean (open dot) and whiskers, representing 1.5x inter quartile range (IQR). Single data points are shown for n≤10. The R² coefficient of determination was determined as: R² = 1 - SSE/SSD (SSE = sum of the squared errors; SSD = sum of the squared deviations about the mean). The linear regressions were estimated with robust standard errors (White heteroscedasticity consistent).

## Extended data

**Extended data Figure 1.**
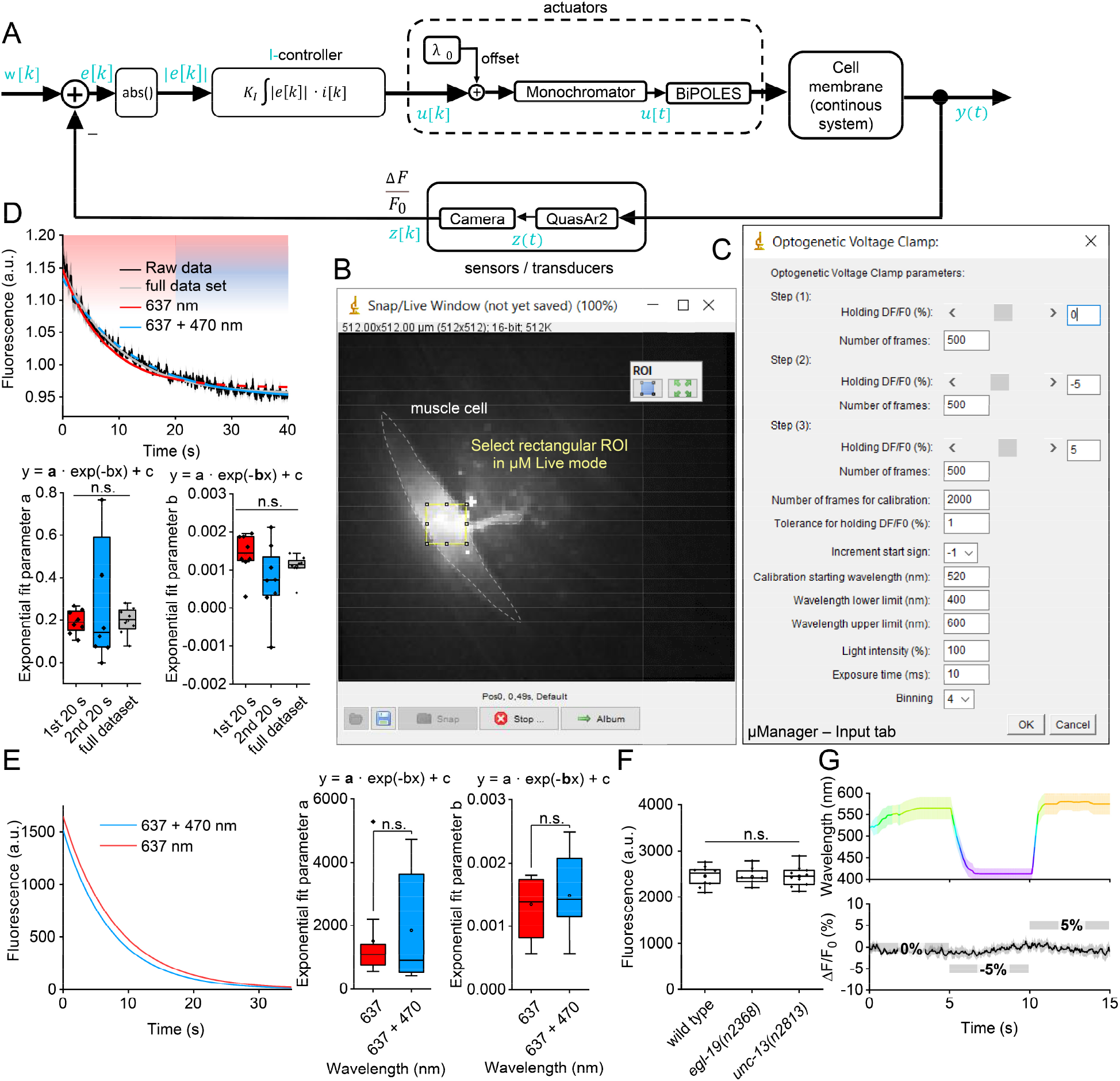
Setup of the OVC software and control parameters. **(A)** Wiring diagram of I-controller: I-controller generates a digital control value u[k] based on error e[k] for the monochromator, which in turn generates analog light output u[t]. BiPOLES modulates membrane voltage y(t) upon analog light activation that in turn defines QuasAr2’s fluorescence z(t) (transduced analog actual value). Fluorescence is monitored via camera and ΔF/F_0_ is calculated (digital actual value). w[k]: Set point. i[k]: current sign of control error e[k]. **(B)** Rectangular ROI selection in μManager “live mode” prior to OVC measurement. **(C)** Input tab of the standard OVC software, implemented in μManager. **(D)** Bleaching behavior and exponential fits of the fluorescence of QuasAr2 in BWMs, in control animals expressing no actuators. Animals were illuminated with the 637 nm laser for the entire duration, and in addition, with 470 nm (300 μW/mm^2^) for the last 20 s (n = 8). Fits were performed for the first (red) or second 20 sec period (blue), or for the entire duration (grey). Lower panel: Statistical analysis of the deduced fit parameters (n = 8). **(E)** Exponential fits of QuasAr2 fluorescence as in (D) but illuminated with either 637 nm laser or 637 nm laser and 470 (300 μW/mm^2^) for the entire duration. Middle and right panels: Statistical analysis of the fit parameters of data in left panel (n = 11-12). (**F**) Expression levels were assessed in wild type animals, and two different mutant strains expressing the OVC components (all from the same integrated transgene, *zxIs139*), using QuasAr fluorescence (n=9-10). Coefficients of variation were 0.08 (wild type), 0.07 (*egl-19*) and 0.09 (*unc-13*). **(G)** Average data for animals expressing only QuasAr2 in BWMs, while the OVC attempts to run a 0, -5, 5 % ΔF/F_0_ protocol (n = 8). Upper panel: Monochromator wavelength; lower panel: mean fluorescence traces. Statistically significant differences analyzed by One-way ANOVA or two-sided t-test with Bonferroni correction (in D, E). Box plots (median, 25-75^th^ quartiles); open dot: mean; whiskers: 1.5x inter-quartile range (IQR).

**Extended data Figure 2.**
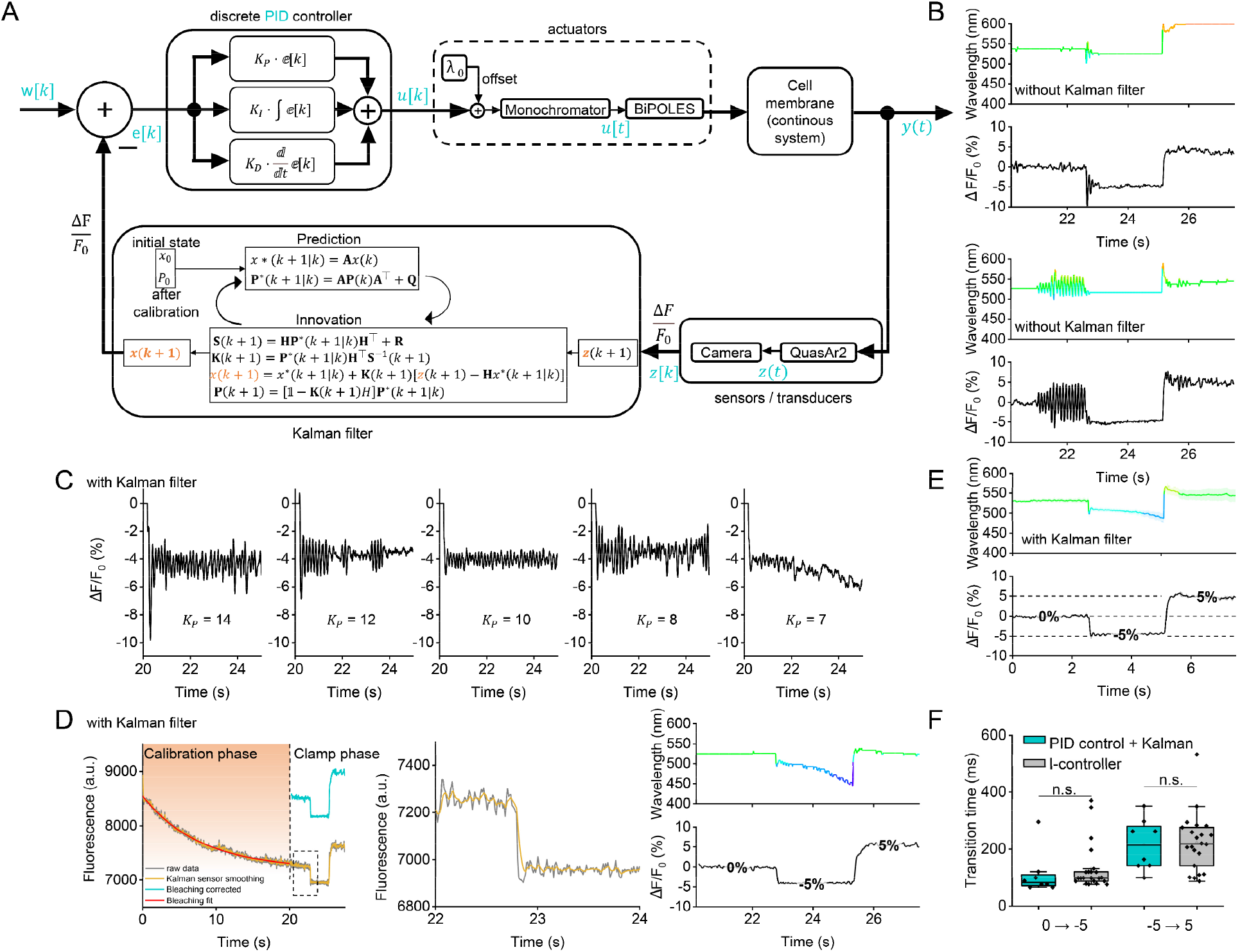
Implementation of a PID controller with Kalman filter. **(A)** Wiring diagram of PID-controller with Kalman filter. Signal flow as in **Extended data Fig. 1A** with the addition of a Kalman filter. **(B)** Comparison of single OVC experiments with PID control but without Kalman filter. Top: No oscillations. Bottom: With strong oscillations, despite identical parameters, emphasizing the need for a Kalman filter to obtain a stable PID-controller. **(C)** Ziegler-Nichols parameter tuning, with Kalman filter. **(D)** OVC three-step protocol (0, -5 and 5 % ΔF/F_0_) in BWMs, insets for close-up. Wavelength shown in the respective color, holding values are indicated for each step, yellow trace represents data processed with Kalman filter for sensor smoothing. Orange shade in left panel: transition period to reach tolerance range. **(E)** Upper panel: Overlay of mean (± S.E.M.) wavelength and (lower panel) fluorescence traces (n=8; holding values: 0, -5, 5 % ΔF/F_0_). **(F)** Times required for the indicated 5 and 10 % ΔF/F_0_ transitions for PID-(plus Kalman filter) and I-controller (n=8-22). Statistically significant differences analyzed by Two-sided t-test with Bonferroni correction. Box plots (median, 25-75^th^ quartiles); open dot: mean; whiskers: 1.5x inter-quartile range (IQR).

**Extended data Figure 3.**
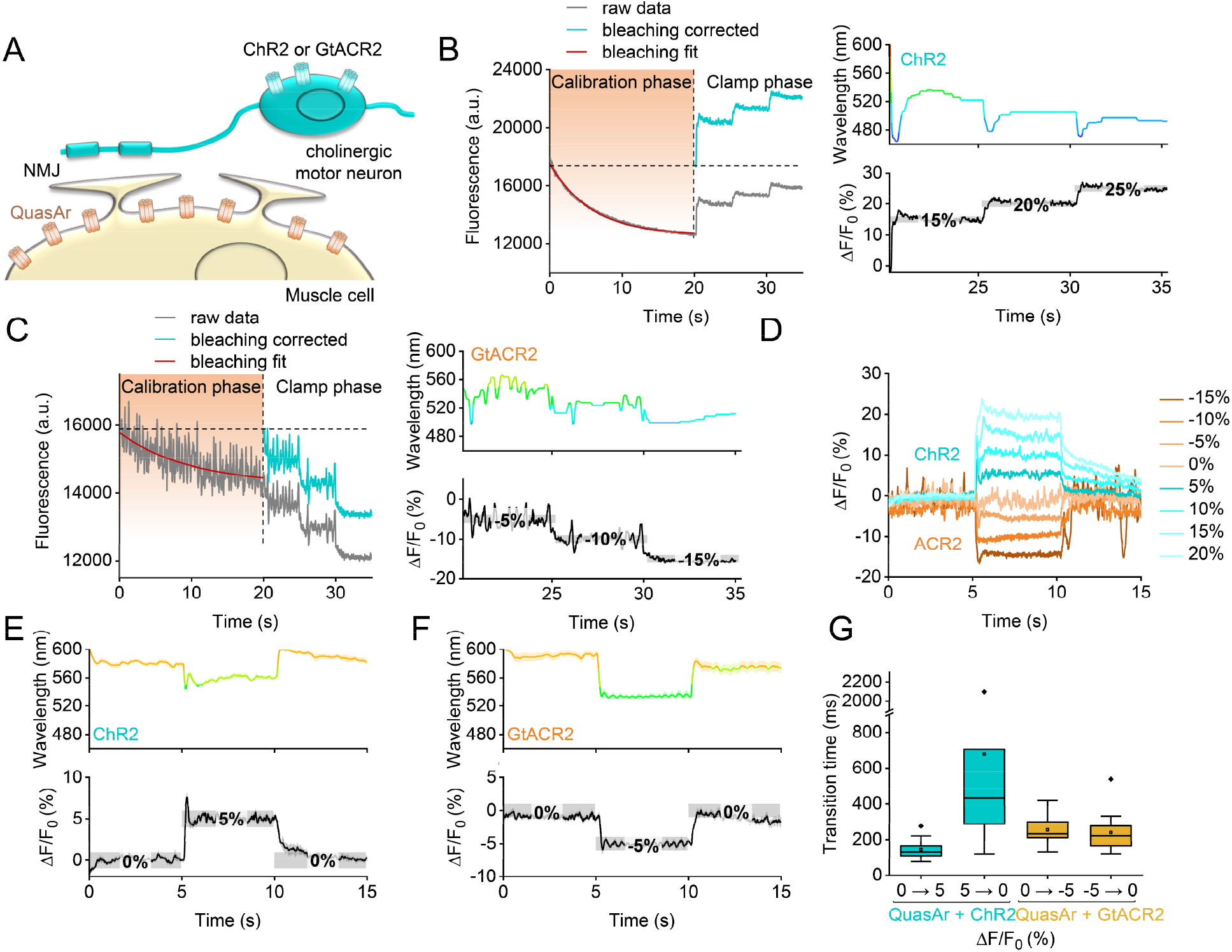
Unidirectional steering of membrane voltage. **(A)** Cholinergic neurons express either ChR2 or *Gt*ACR2, while QuasAr2 is expressed in BWMs. **(B)** Three-step protocol in ChR2 animals. Left: Raw and bleaching corrected data of calibration (red: exponential fit) and clamping phase. Right panels: Corresponding wavelength and ΔF/F_0_ traces. Holding values 15, 20 and 25 % ΔF/F_0_; grey shade: tolerance range for each step. **(C)** Three-step (−5, -10 and -15 % ΔF/F_0_) protocol in *Gt*ACR2 animals, as in (B). **(D)** Single traces for both strains, 0 %, then -15 to 20 % ΔF/F_0_ (in 5% increments), and return to baseline. **(E)** Mean (± S.E.M.) traces (n = 17); holding values: 0, 5, 0 % ΔF/F_0_, in ChR2 animals. **(F)** As in E, for *Gt*ACR2 animals (n = 14; 0, -5, 0 % ΔF/F_0_). **(G)** Transition time, respective 5 % ΔF/F_0_ steps; box plots (median, 25-75^th^ quartiles); open dot: mean; whiskers: 1.5x inter-quartile range (IQR).

**Extended data Figure 4.**
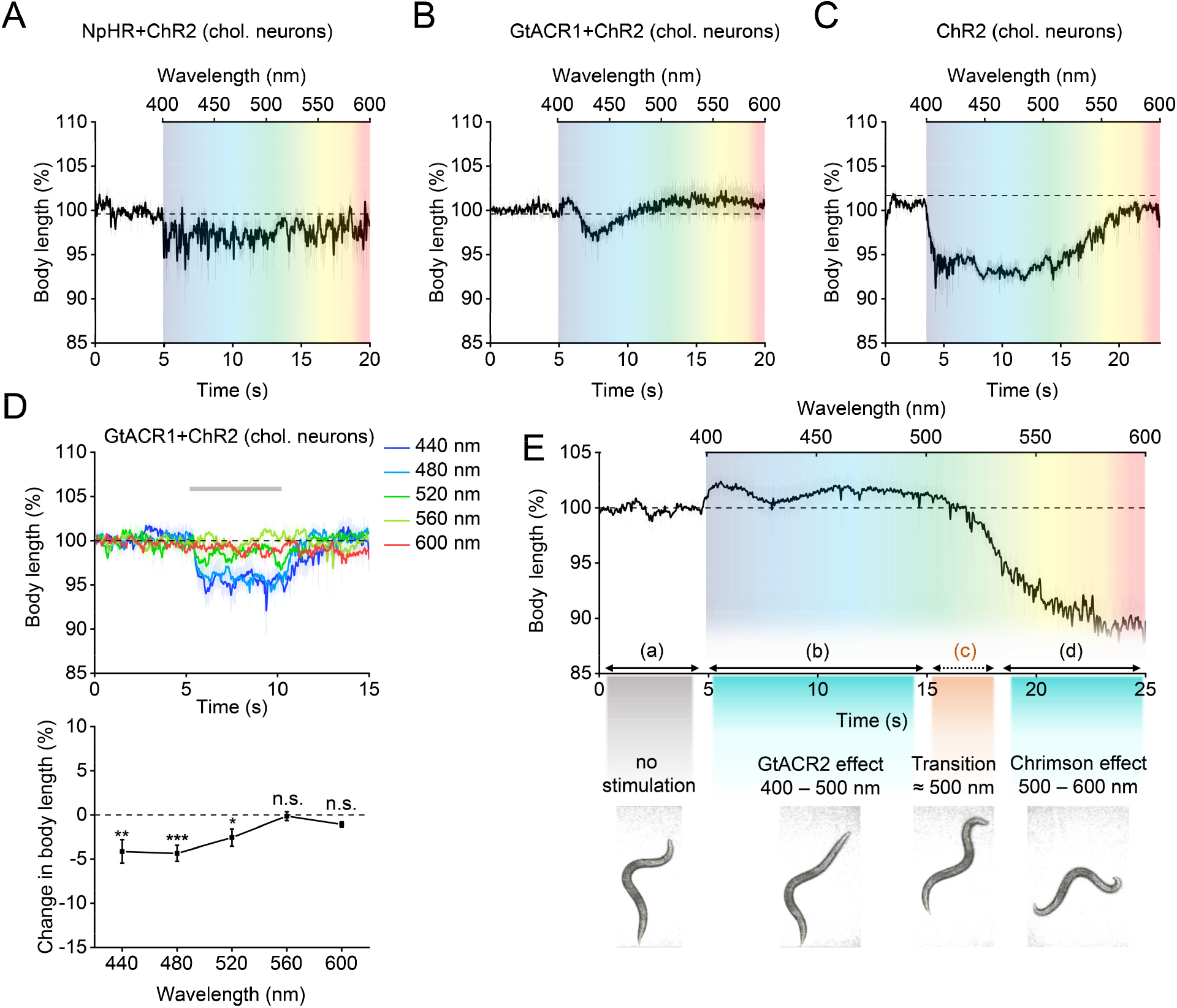
Testing different combinations of depolarizing and hyperpolarizing actuators. **(A-E)** Body length measurements to test functionality of optogenetic actuator combinations, expressed in cholinergic neurons. (A) NpHR and ChR2 (n = 8, monochromator wavelength ramp from 400 to 600 nm, 300 μW/mm^2^). (B) *Gt*ACR1 and ChR2 (n = 10). (C) ChR2-only (n = 8). (D) *Gt*ACR1 and ChR2. Upper panel: 5 s light pulses, wavelength as indicated, 300 μW/mm^2^ (n = 5-7). Lower panel: Mean (± S.E.M.) body length changes. Statistically significant differences were analyzed by one-way ANOVA with Bonferroni correction (***P ≤ 0.001, **P ≤ 0.01, *P ≤ 0.05). (E) Upper panel: Body length measurements of animals expressing BiPOLES in cholinergic neurons (mean ± S.E.M., wavelength ramp 400 - 600 nm, n = 8). Below: Representative still images of animals for each phase of the experiment: (a) before light, (b) *Gt*ACR2 effect / muscle relaxation, (c) transition from hyper- to depolarization, (d) Chrimson effect / muscle contraction.

**Extended data Figure 5.**
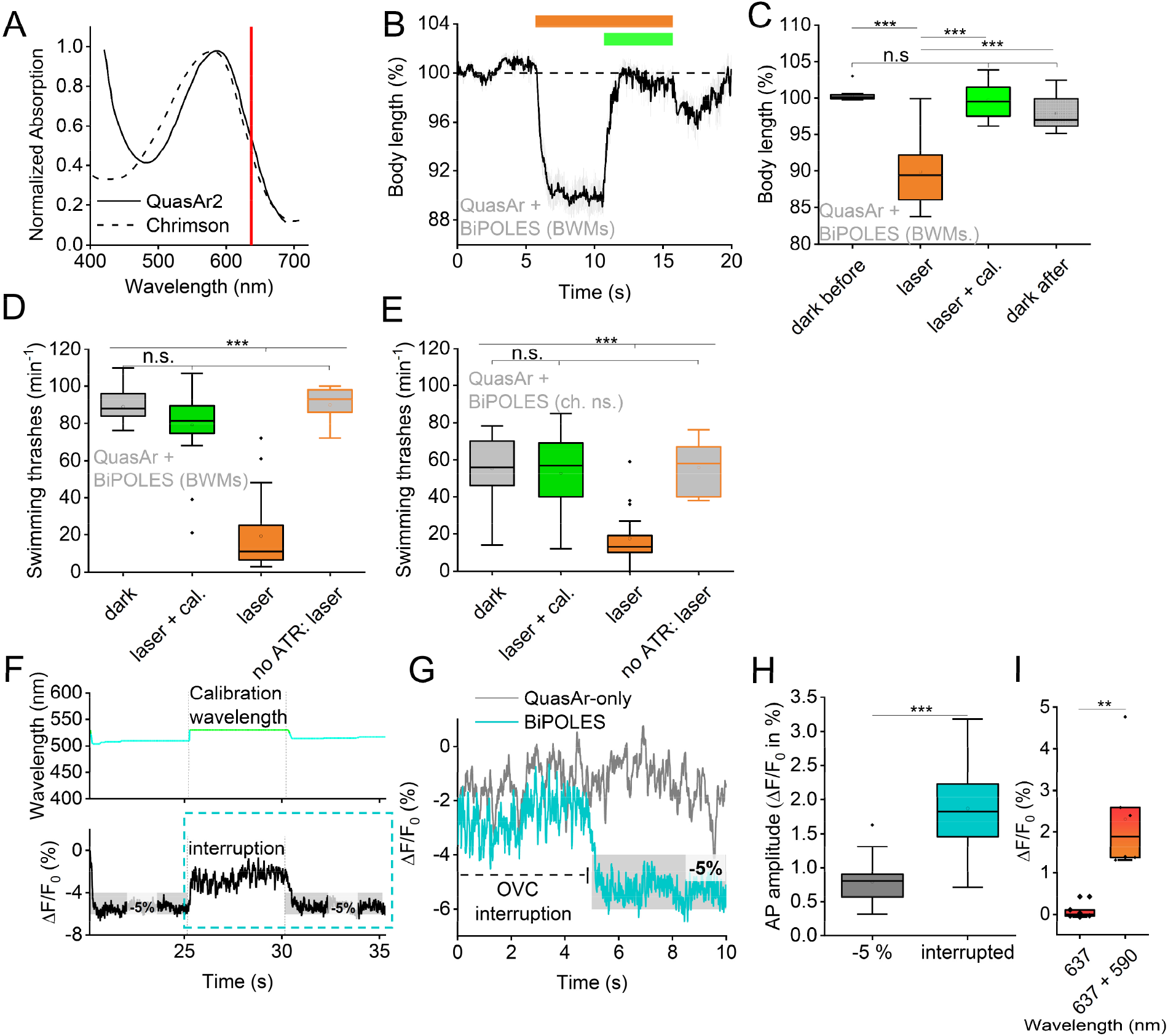
BiPOLES activation by 637 nm laser light and calibration wavelength have no adverse effects on muscle function and locomotion. **(A)** Normalized absorption spectra of Chrimson and QuasAr2^26,39^, laser wavelength used for QuasAr2 imaging is indicated by a red line. **(B)** Body length measurements of animals expressing BiPOLES and Quasar in BWMs (mean ± S.E.M., light pulses: 637 nm laser and 637 nm laser plus calibration wavelength (ca. 521 nm, n = 14). **(C)** Statistical analysis of data in (B). **(D, E)** Analysis of swimming activity of animals expressing BiPOLES and Quasar in BWMs (D, n = 9-20) or cholinergic neurons (E, n = 10-18). Illumination parameters as in (C), or in the absence of all-*trans* retinal (ATR). **(F)** OVC experiment in BWMs (BiPOLES and QuasAr2) with clamp interruption (protocol: 1) holding value - 5 %, wavelength determined by OVC, 2) interruption at calibration wavelength, 3) holding value - 5 % ΔF/F_0_). **(G)** Close-up of (F, lower panel), with comparison to voltage fluorescence activity of unstimulated animals expressing only QuasAr2. **(H)** Group data of ΔF/F_0_ fluorescence amplitudes of typical, action potential-based fluctuations during -5 % clamping, or during no OVC action (n=30). Box plots (median, 25-75^th^ quartiles); open dot: mean; whiskers: 1.5x inter-quartile range (IQR). **(I)** Mean ΔF/F_0_ QuasAr2 signal (n=6) in BWMs (co-expressed with BiPOLES) in the presence of laser light (637 nm) and additional signal in response to Chrimson excitation light (590 nm). In (C-E, H, I): Statistically significant differences were analyzed by two-sided t-test or One-way ANOVA with Bonferroni correction (***P ≤ 0.001, **P ≤ 0.01, *P ≤ 0.05). Box plots (median, 25-75^th^ quartiles); open dot: mean; whiskers: 1.5x inter-quartile range (IQR).

**Extended data Figure 6.**
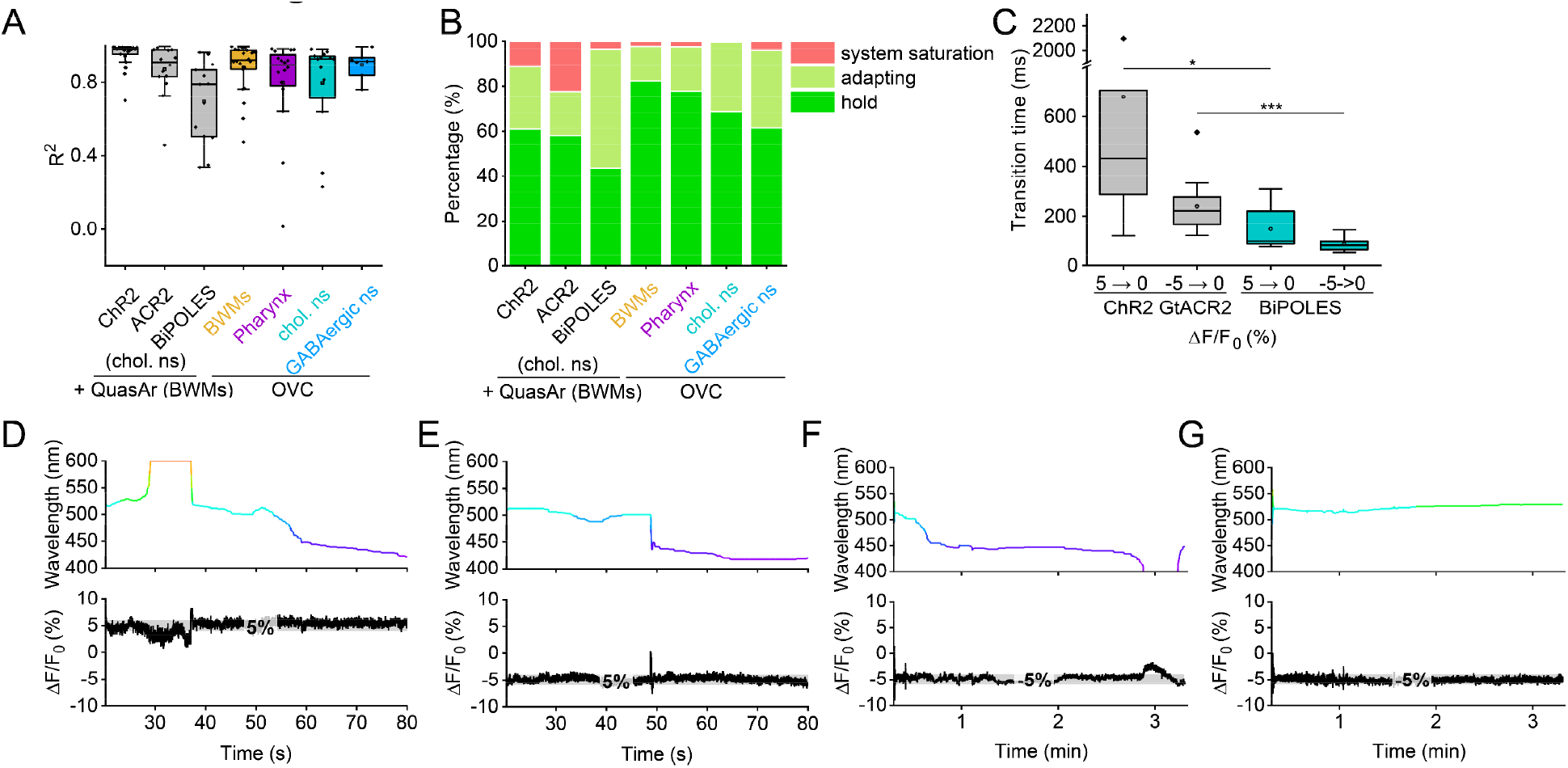
Performance assessment, and long-term action of the OVC. **(A)** Quality of the exponential fit for bleach correction (coefficient of determination, R^2^), for the different QuasAr2 / optogenetic actuator combinations, as indicated (n=7-22). **(B)** Percentages of the clamping status (hold, adapting, system saturation), for the different QuasAr2 / optogenetic actuator combinations, as indicated (n=7-22). **(C)** Summary of OVC speed, reflected by the time needed for a 5 % ΔF/F_0_ step for different configurations (single tools, same-cell approach; n=10-22). **(D-G)** Long-term OVC action, de- and hyperpolarizing steps. Shown are the achieved fluorescence values (lower panels, grey shades are tolerance range of the OVC protocol), and monochromator wavelength required (upper panels). Statistically significant differences were analyzed (C) by one-way ANOVA with Bonferroni correction (***P ≤ 0.001, *P ≤ 0.05).

**Extended data Figure 7.**
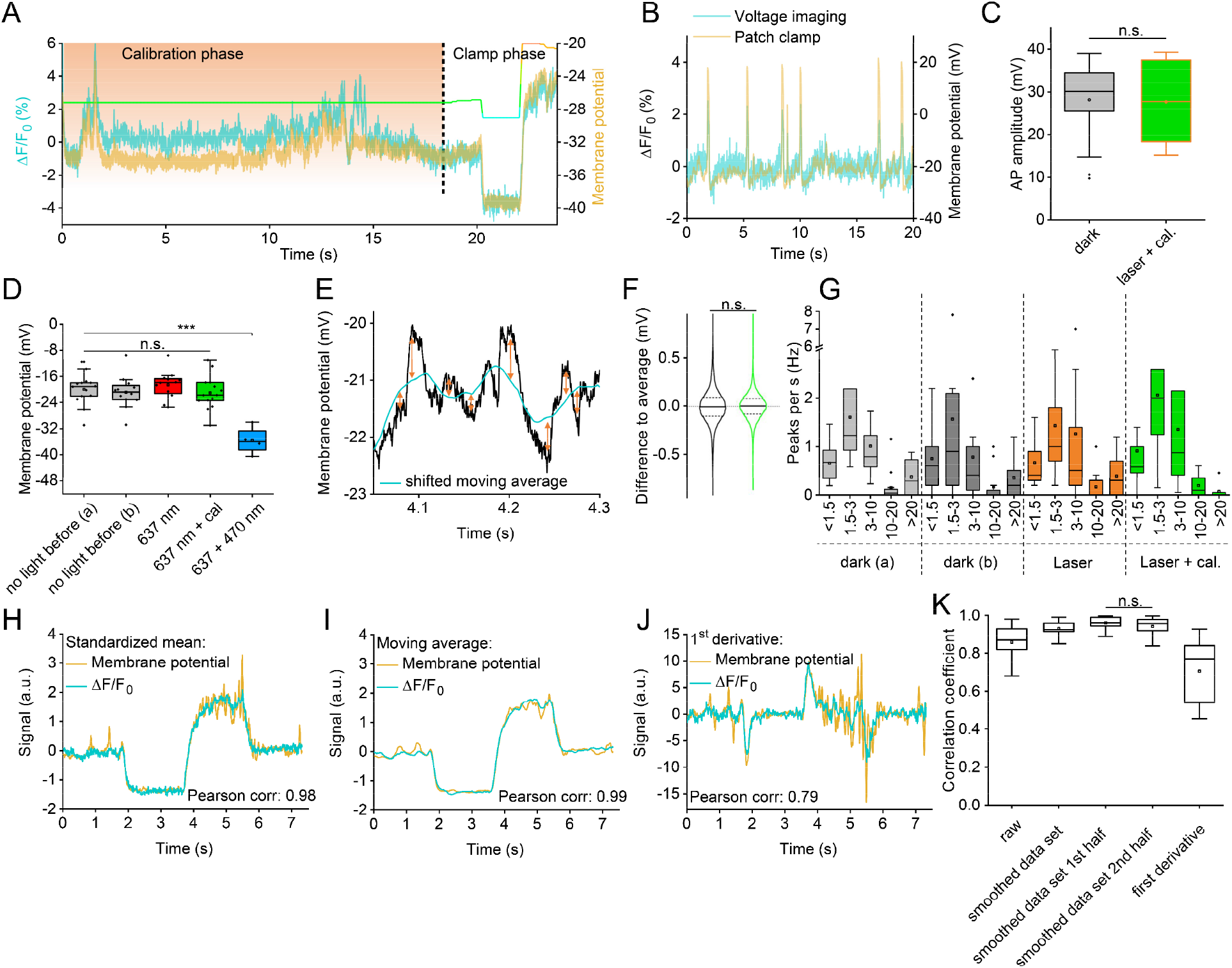
Simultaneous patch-clamp and fluorescence measurement, normal membrane voltage behavior in BiPOLES-activated BWMs, no progressive error following calibration phase. **(A)** Original record of simultaneous voltage and fluorescence measurement during calibration and clamp phase (3 step OVC protocol, -3, +3 % ΔF/F_0_). Note the fluorescence trace was subsequently bleaching-corrected for the calibration phase. **(B)** APs in simultaneous patch-clamp and fluorescence recordings during OVC calibration phase (637 nm laser and 521 nm calibration wavelength). **(C)** Statistical analysis of AP amplitude. n = 4, 24-27 APs. **(D)** Membrane potential in muscle was measured by patch-clamp under the indicated light conditions. **(E, F)** Analysis of small, subthreshold voltage fluctuations observed during patch-clamp, without or with 637 nm laser and calibration wavelength. Differences (arrows) of actual peaks to shifted moving average (blue), as a proxy for base line (E), were statistically analyzed in (F). n=14, with ca. 55.000 single data points per measurement. **(G)** Frequency distribution of distinct voltage signals, incl. APs, observed during patch-clamp recordings. Mean number of events (±S.E.M.) per second is shown as a function of peak amplitudes (in mV), for the different illumination conditions, as indicated below. n = 14, 3118 peaks. None of the respective amplitude distributions showed significant differences to any other bin. **(H)** Patch-clamp derived membrane voltages (mean, yellow line) were compared to the mean ΔF/F_0_ levels (blue line) induced by the OVC during the clamp phase, based on parameters derived in the calibration phase (not shown). **(I)** As in (H), but mean moving averages were analyzed. **(J)** As in (H), but the signal change (1^st^ derivative) was compared. **(K)** Statistical analyses of correlation coefficients determined from data in (H-J), comparing the first and second halves of the experiments (p = 0.31). Correlation coefficients show high fidelity, typically >0.8. Statistically significant differences (***P ≤ 0.001) were analyzed by one-way ANOVA with Bonferroni correction (in F, G, K) or paired, two-sided t-test (in C). Box plots (median, 25-75^th^ quartiles); open dot: mean; whiskers: 1.5x inter-quartile range (IQR).

**Extended data Fig. 8.**
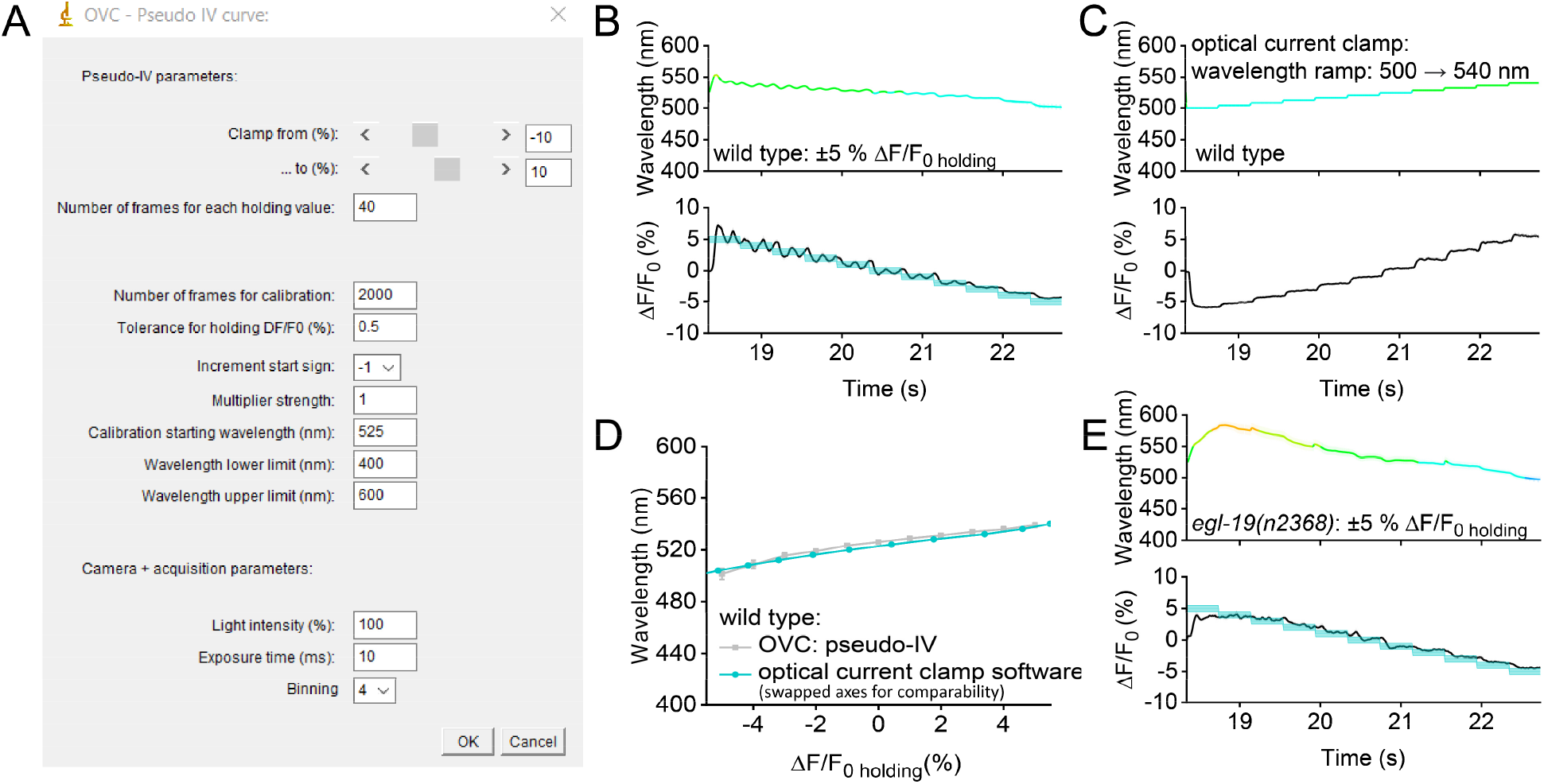
Optical pseudo-I/V curve measurements: **(A)** Input tab of the “pseudo I/V curve” software allowing to run distinct % ΔF/F_0_ as clamp values (equivalent to voltages), while recording wavelengths (equivalent to currents). **(B)** Ramping +5 to -5 % ΔF/F_0_ (lower panel, mean ±S.E.M.; blue shades: tolerance ranges), while recording wavelengths (upper panel, mean ±S.E.M.; n=17). **(C)** Inverse experiment of (B), single trace, running a wavelength ramp (upper panel) and recording % ΔF/F_0_ (lower panel; n=15). **(D)** Comparison of OVC-based pseudo-I/V curve and measurement with the optical current clamp software (see **Extended data Fig. 9A-C**), demonstrating high fidelity of the OVC control capabilities. **(E)** Measuring optical pseudo-I/V curves for *egl-19(n2368)* mutants, compare to (B) for wild type animals (n=14).

**Extended data Fig. 9.**
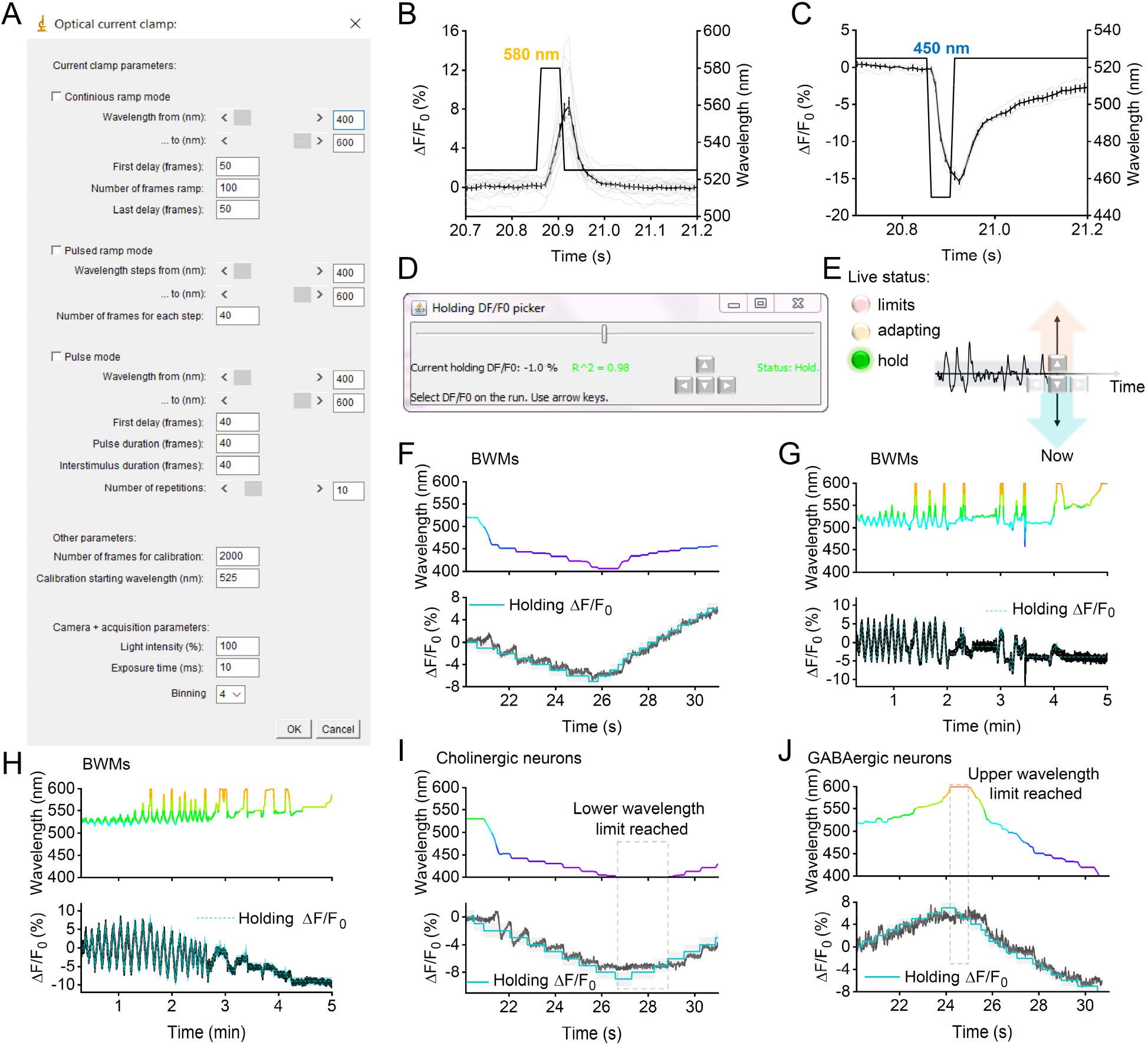
Software for ‘optical current clamp’, and software for ‘on-the-run’ live voltage adjustment: **(A)** Software to achieve bidirectional optical current clamping, input tab. **(B, C)** Mean ±S.E.M. % ΔF/F_0_, resulting from 100 ms depolarizing (590 nm) step (B, n=13) or from a hyperpolarizing (450 nm) step (C, n=5). **(D-J)** Time-varying OVC live control (‘on-the-run’). (D) User interface for software version allowing live control of membrane voltage fluorescence. (E) Scheme: Holding values can be selected using arrow keys. Live status (system on hold, adapting or exceeding limits) is shown, enabling adjustment. (F-H) Example traces of wavelength (upper panels) and holding values as % ΔF/F_0_ (lower panels) in BWMs for brief (F) and extended periods (G, H), as well as in cholinergic (I) and GABAergic neurons (J).

**Extended data Figure 10.**
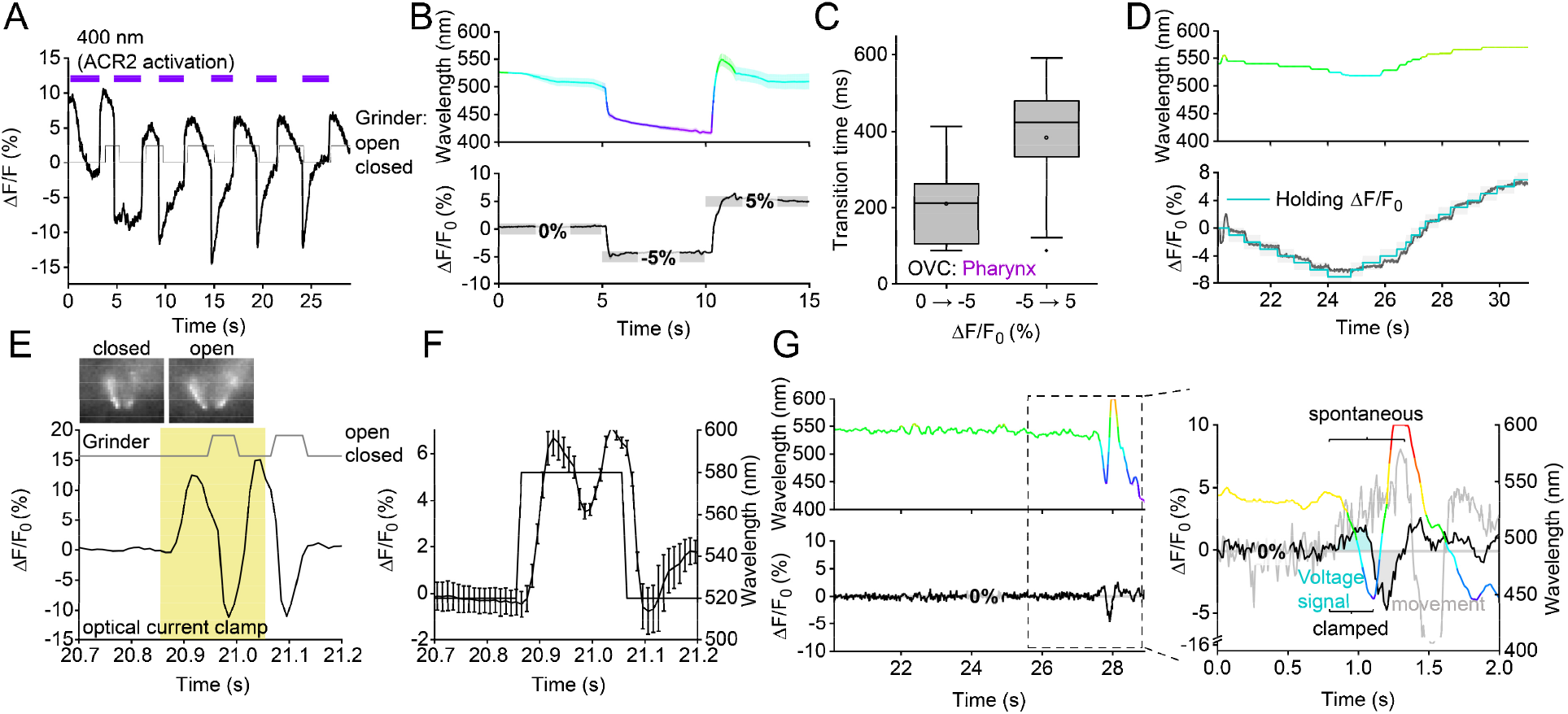
Voltage imaging and OVC measurements in pharyngeal muscle and the motor neuron DVB. **(A)** Voltage imaging and analysis of grinder opening in animals expressing QuasAr2 and BiPOLES in pharyngeal muscle, stimulated with consecutive 400 nm light pulses (300 μW/mm^2^). Corresponding open or closed state of the grinder as indicated. **(B)** Transition time required by the OVC in the pharynx to execute 5 and 10 % ΔF/F_0_ steps. **(C)** Mean (± S.E.M.) traces (n = 16) of OVC protocol in pharyngeal muscle (0, -5, 5 % ΔF/F0). **(D)** “On-the-run”-experiment of OVC in pharyngeal muscle. **(E, F)** ‘Optical current clamp’ protocol, applied to pharyngeal muscle (yellow shade, depolarization evokes two APs), and assessing QuasAr fluorescence as a readout; single experiment (E), mean fluorescence analysis (n=9, F). **(G)** Example DVB voltage fluorescence (lower panel) and wavelength (upper panel) traces upon suppression of an AP by the OVC. Inset: Close-up and overlay of spontaneous (light gray trace) and clamped (black trace) DVB voltage signal. Monochromator wavelength is presented in the respective color. Voltage signal and movement artefact highlighted in blue and grey, respectively.

**Extended data video 1. *C. elegans* expressing BiPOLES in cholinergic motor neurons, during a wavelength ramp from 400-600 nm**. Animal crawling on solid substrate, with inactive (relaxed) muscle, transition to normal movement, followed by activated (contracted) muscle and respective paralysis, as indicated.

**Extended data video 2: ‘On-the-run’ mode of the OVC, enabling adjusting membrane voltage during a running acquisition**. Left panels, upper: monochromator wavelength, lower: voltage fluorescence trace. Arrow keys indicate the live settings chosen by the experimenter.

**Extended data video 3: Dynamically clamping voltage in the pharynx, using the OVC**. Video shows QuasAr fluorescence in the terminal bulb of the pharynx. Structures of the grinder of the pharynx are indicated (open and closed states), as well as the calibration and clamping phase. Overlaid are optical voltage traces (bottom), as well as the wavelength of the monochromator, used during clamping, as a color-coded trace at the top.

## Notes

### Summary of Updates

The revised manuscript now includes a more elaborate calibration and new software that allows to generate optical pseudo-IV curves, directly translating fluorescence signals to membrane potential and wavelengths to currents (Figure 2). This way, we were able to analyze the I/V-relationship of the mutated channel in *egl-19* VGCC gain-of-function mutants (Figure 3). In addition, we now provide simultaneous behavioral analysis of the OVC pharynx AP suppression, where the system could not only dynamically follow and counteract native APs, but also fully suppress associated behaviors (Figure 6, E.D. video 3). We also demonstrate long-term OVC application of up to 5 minutes (E.D. Figure 9 G, H). Besides our all-optical voltage clamp approach, a further software now provides an optical adaptation of a current clamp, by live recording of QuasAr's fluorescence and presenting either single light pulses at a desired wavelength or stepwise or continuous wavelength ramps (E.D. Figure 9 A-C, E.D. Figure 10 E, F).

